# Asterix/Gtsf1 links tRNAs and piRNA silencing of retrotransposons

**DOI:** 10.1101/2020.08.11.246769

**Authors:** Jonathan J. Ipsaro, Paul A. O’Brien, Shibani Bhattacharya, Arthur G. Palmer, Leemor Joshua-Tor

## Abstract

The piRNA pathway safeguards genomic integrity by silencing transposable elements in the germline. While Piwi is the central piRNA factor, others including Asterix/Gtsf1 have also been demonstrated to be critical for effective silencing. Here, using eCLIP with a custom informatic pipeline, we show that Asterix/Gtsf1 specifically binds tRNAs in cellular contexts. We determined the structure of mouse Gtsf1 by NMR spectroscopy and identified the RNA binding interface on the protein’s first zinc finger, which was corroborated by biochemical analysis as well as cryo-EM structures of Gtsf1 in complex with co-purifying tRNA. We further show that LTR retrotransposons are preferentially de-repressed in Asterix mutants. Given the role of tRNAs as LTR retrotransposon primers, our work implicates Asterix/Gtsf1 as exploiting tRNA dependence to identify transposon transcripts and promote piRNA silencing.

## Introduction

To maintain genomic integrity, the activity of mobile genetic elements (transposons) must be repressed. This is particularly important in the germline where transposon silencing, enforced by the Piwi-interacting RNA (piRNA) pathway^1,2^, affords genetic stability between generations. piRNA silencing is accomplished through interrelated mechanisms that function in distinct cellular compartments. In the cytoplasm, piRNA-directed cleavage leads to post-transcriptional target degradation^3,4^. In the nucleus, however, Piwi-piRNA complexes are believed to recognize nascent transposon transcripts, recruit additional factors, and ultimately enforce the deposition of histone H3 lysine 9 trimethylation (H3K9me3) repressive marks^5–8^.

The results of three, independent, genome-wide screens revealed a number of candidate proteins that are essential for piRNA silencing^9–11^. While many of these factors affected piRNA biogenesis, others had no obvious effect on piRNA levels or composition. This suggests that this second class of factors principally act downstream of piRNA biogenesis most likely either in transcriptional gene silencing (TGS) or in the ping-pong cycle (reviewed in Czech *et al.*^1^).

Recent work on several of these proteins (including, but not limited to, Panoramix^12^, Nxf2, and Nxt1^13–15^) has provided a framework for linking the piRNA pathway to deposition of heterochromatic silencing marks. However, many of the molecular and mechanistic underpinnings that govern these connections remain obscure. With this in mind, we endeavored to detail the role of one of the strongest hits in the aforementioned screens, the protein CG3893/Cue110/Asterix/Gtsf1, in piRNA transposon silencing.

In *Drosophila*, expression of Asterix is largely restricted to the female germline, where it is critical not only to transposon silencing, but also more broadly for ovarian development. There, Asterix localizes to the nucleus and has been shown to interact with Piwi^11,16,17^. Similarly, gametocyte-specific factor 1 (Gtsf1), the mammalian homolog of Asterix, is involved in retrotransposon suppression and is also important in both oogenesis and spermatogenesis^18,19^. Reports on the sub-cellular localization of Gtsf1 are mixed, with the most recent findings revealing focal localization in both nuclei and cytoplasmic processing bodies (piP bodies)^19^.

Asterix and Gtsf1 are small proteins, 167 amino acids in length, predicted to consist of two N-terminal CHHC-type zinc fingers and a disordered C-terminal domain (Fig. 1a). CHHC zinc fingers have only been identified in eukaryotes and there are found in just three protein groups: spliceosomal U11-48K proteins, tRNA methyltransferases, and gametocyte specific factors (such as Asterix/Gtsf1)^20^. In the former two cases, these motifs have been demonstrated to bind RNA^21,22^.

**Figure 1.**
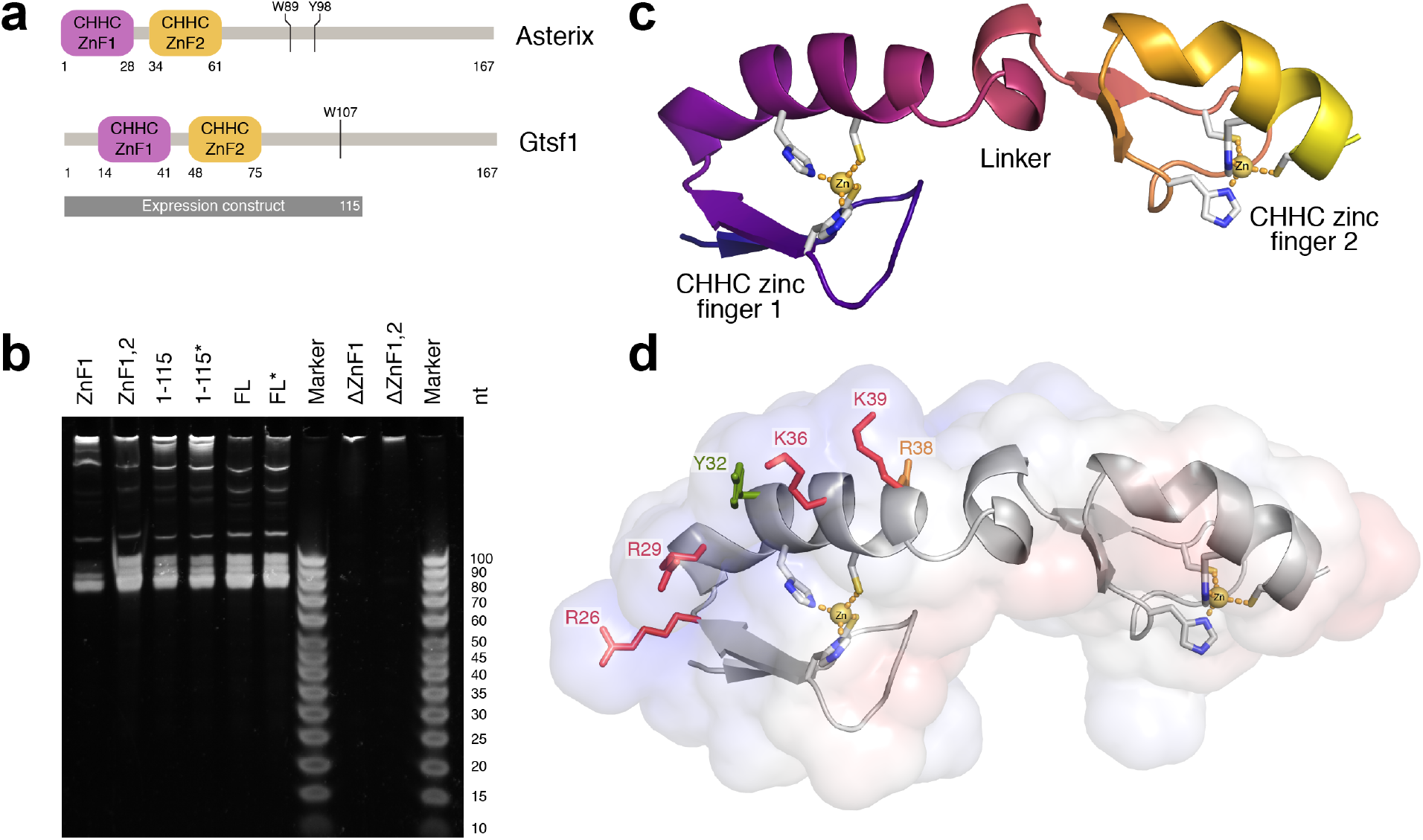
Structure and RNA binding activity of Asterix/Gtsf1. **(a) Domain architecture.** Asterix/Gtsf1 is comprised of two, N-terminal, CHHC zinc fingers and a C-terminal tail predicted to be disordered. Aromatic residues that interact with Piwi proteins are indicated. **(b) Urea-PAGE analysis of RNAs that co-purify with Gtsf1 truncation constructs.** Domains or amino acid ranges of each recombinantly-expressed mouse Gtsf1 construct are indicated above the corresponding lane. FL denotes full-length protein. Asterisks indicate constructs containing four cysteine to serine point mutations that were included in the NMR construct to limit aggregation. **(c) Solution structure of mouse Gtsf1.** The lowest energy structure for the protein’s core (residues 13-72) is depicted as a ribbon diagram. Zinc-coordinating residues are shown as sticks with zinc atoms displayed as yellow spheres. **(d) Mapping of the RNA-binding interface.** The calculated electrostatic surface of mouse Gtsf1 (scaled from −5 k_B_T in red to +5 k_B_T in blue) displays a positively-charged ridge on ZnF1. Zinc-coordinating residues and point mutations tested for effects on RNA binding are shown as sticks (red = abolishes binding; orange = hinders binding; green = no effect).

To detail the role of Asterix/Gtsf1 in retrotransposon silencing, we implemented a combination of biochemical, structural, cell-based, and informatic analyses. Here, we present biochemical evidence that Asterix/Gtsf1 directly binds RNA. We determined the structure of mouse Gtsf1 using NMR spectroscopy and mapped the RNA binding site through mutational analysis. Using eCLIP and a customized informatic workflow, we demonstrate that Asterix/Gtsf1 preferentially binds tRNAs in cells. Using cryo-electron microscopy we solved a low-resolution structure of Gtsf1 in complex with tRNA. Together, these findings led us to propose a model of how Asterix uses tRNA biology to effect transposon silencing. Informatic analysis of existing datasets implicated Asterix as particularly relevant in silencing LTR transposons, a transposon class which share an evolutionary history with tRNA.

## Results

### Asterix/Gtsf1 is an RNA-binding protein

We initiated structural studies with recombinantly-produced mouse Gtsf1 to systematically characterize its molecular role in retrotransposon silencing. During purification from *Sf9* cells, we observed that Gtsf1 co-purified with endogenous nucleic acids (Supplemental Fig. 1). These species could be separated by ion exchange chromatography (Supplemental Fig. 1a), resulting in monodisperse and highly-purified protein (Supplemental Fig. 1b-c) in addition to isolated nucleic acids.

When assessed by urea-PAGE (Supplemental Fig. 1d), the nucleic acids displayed a narrow size distribution (~70-90 nucleotides), suggesting that specific ligands were being bound. We hypothesized that this Gtsf1-bound material was RNA by analogy to other CHHC zinc fingers proteins’ ligands. Treatment with RNase A or sodium hydroxide degraded this material whereas treatment with DNase I did not, verifying that this was indeed the case (Supplemental Fig. 1e).

To further pinpoint the RNA binding activity of Gtsf1, we created a panel of truncation constructs, similarly expressed each in *Sf9*, and assessed which of these co-purified with RNA (Fig. 1b). The first CHHC zinc finger (ZnF1) was found to be both necessary and sufficient for the majority of RNA binding (Fig. 1b). Additional inclusion of the second CHHC zinc finger fully recapitulated the RNA size profile as compared of the full-length protein’s pulldown.

Given that purified RNAs are usually unstable and RNAs of this size are unlikely to be fully protected by a single, 45 amino-acid (~5 kilodalton) protein binding partner such as ZnF1, this result also suggesting that the isolated RNAs were structured, affording them some protection from degradation.

### Overall structure of Gtsf1

As the zinc finger RNA-binding modules were now of primary interest, we examined a construct of mouse Gtsf1, spanning residues 1 to 115, using NMR spectroscopy (Fig. 1c, Supplemental Figs. 2-3, Supplemental Table 1). In agreement with folding and domain predictions, initial HSQC experiments suggested the protein to contain both ordered and disordered segments (Supplemental Fig. 2a). Subsequent backbone assignment more specifically indicated the structured core of the protein spanned residues 13-73, with residues outside this range tending to be disordered. Analysis of secondary chemical shifts (Supplemental Fig. 2d-e) as well as backbone conformation predictions using TALOS and CSI methods indicated strand-strand-helix architectures for both ZnF1 and ZnF2, similar to that observed for the only other reported CHHC zinc finger structure^21^.

Structure determination of residues 1-80 revealed two, tandem, CHHC zinc finger domains (ZnF1, ZnF2) connected by an α-helix-containing linker (Fig. 1c, Supplemental Fig. 3) with the N- and C-termini being extensively disordered. In preliminary structure calculations which did not include restraints for the CHHC residues with zinc, each zinc finger nevertheless adopted a strand-strand-helix fold with the appropriate zinc-coordinating residues in proximity to one another.

In the NMR-derived structural ensemble of the twenty, final, lowest energy structures (Supplemental Fig. 3a-c), the relative positions of the zinc finger domains varied somewhat owing to flexibility in the intervening linker. Nonetheless, structure calculations for the individual domains were highly superimposable (Supplemental Fig. 3f-g) with R.M.S.D. values of 0.2 Å for backbone atoms for each zinc finger. Moreover, these domains were superimposable with each other and the only other CHHC zinc finger structure available (from the U11-48K spliceosomal protein^21^) (Supplemental Fig. 3h). Co-evolution analysis^23^ additionally corroborated the overall protein fold, with several intra-ZnF residues displaying evidence of co-evolution (Supplemental Fig. 3e). Final validation of the structural ensemble with Molprobity^24^ indicated reasonable geometry overall, with the core (residues 13-73) possessing very few violations (Supplemental Table 1).

### ZnF1 presents a conserved RNA-binding interface

Guided by the protein structure, we next mapped the RNA-binding interface. Calculation of the electrostatic surface of Gtsf1 revealed a pronounced, positively-charged ridge running the length of ZnF1 (Fig. 1d). Mutagenesis of single basic residues along this patch abrogated or reduced RNA-binding activity with no apparent effects on expression or solubility (Supplemental Fig. 4a,c), indicating that they indeed form part of the RNA-binding interface.

Evolutionary analysis corroborated the importance of ZnF1, with residues important for RNA binding among the most highly conserved in the structure (Supplemental Fig. 3d). Although some key residues—notably in the CHHC metal-coordination site—of ZnF2 were also highly conserved, ZnF2 was more variable overall. Correspondingly, mutations of basic residues on ZnF2 (Supplemental Fig. 4b,d) did not affect RNA-binding.

Together, these findings bolstered our initial characterization that ZnF1 mediates RNA interactions (Fig. 1b) and precisely identified basic residues in this region as forming a conserved interface for RNA-binding.

### Recombinantly-produced Gtsf1 co-purifies with tRNAs

To complement the biochemical characterization of Gtsf1 protein and gain insight into the possible identities of biologically-relevant ligands, we next analysed the RNAs that were being retained during recombinant expression in *Sf9*. RNAs that co-purified with mouse Gtsf1 were isolated by phenol:cholorform extraction, ethanol precipitated, then subjected to size-selection and next-generation sequencing.

Consistent with the previous observation that the bound RNAs were approximately 70-90 nucleotides in size, we found considerable enrichment of tRNAs in the Gtsf1 pull-down compared to size-matched controls (Supplemental Table 2). This enrichment was readily apparent, even though the *Sf9* genome is not fully annotated, as approximately 15% of the sequencing library was comprised of a single tRNA species. Moreover, each of the twenty most abundant sequences were determined to be tRNA-derived, with 50% of all library reads corresponding to these twenty sequences.

This contrasted with size-matched controls from extracted *Sf9* total RNAs where the top sequence was derived from the highly abundant large ribosomal subunit yet nonetheless made up only ~4% of the library. The top tRNA read in the size-matched control only contributed approximately 0.3% of the total reads.

### Gtsf1 directly binds tRNAs in cellular contexts

To catalogue RNAs interacting directly with Gtsf1 in a cellular context, we employed eCLIP^25^. Strep-tagged Gtsf1 was transfected into a mouse embryonal teratoma cell line (P19), bound RNAs were covalently linked using UV-crosslinking, the complexes isolated by affinity purification, and the RNA subjected to next-generation sequencing.

Many classes of RNA—such as rRNAs, tRNAs, and highly repetitive genetic elements like transposons—are typically excluded from downstream analysis due to ambiguity in read mapping and/or their high abundance. Given the relevance of these gene classes in the context of Asterix/Gtsf1, we therefore developed a custom bioinformatic workflow to ensure their inclusion. Read mapping was performed allowing for multimapping with up to 50 genomic sites per read^26^. Various sources of well-curated gene annotations (including gencode^27^, miRBase^28^, tRNAdb^29^, piRNAclusterdb^30^, and TEtranscripts^31^) were integrated for customized annotation calling. We sequentially filtered each read into a single annotation class based on known biological abundance of that class. Subsequently, the enrichment for a given gene or annotation class was determined, first by subtracting levels in a background (non-crosslinked) pull-down, then by comparing to input read counts, with multi-mapping reads being 1/n-normalized across genes within the assigned annotation class.

Analysis of these data by annotation category displayed a substantial enrichment of tRNAs (Fig. 2a) and when analysed as a distribution of fold enrichments for individual genes in each annotation class (i.e. per locus), we again found a preponderance of tRNA enrichment (Fig. 2b). A more granular inspection of the tRNA reads revealed widespread coverage across the tRNA body, suggesting that full-length (or nearly full-length) tRNAs were being bound. Although some variability was observed in the amount of enrichment across different tRNAs, in contrast with the *Sf9* pull-downs, no particular tRNA or set of tRNAs was selected preferentially based on the anticodon (Fig. 2c).

**Figure 2.**
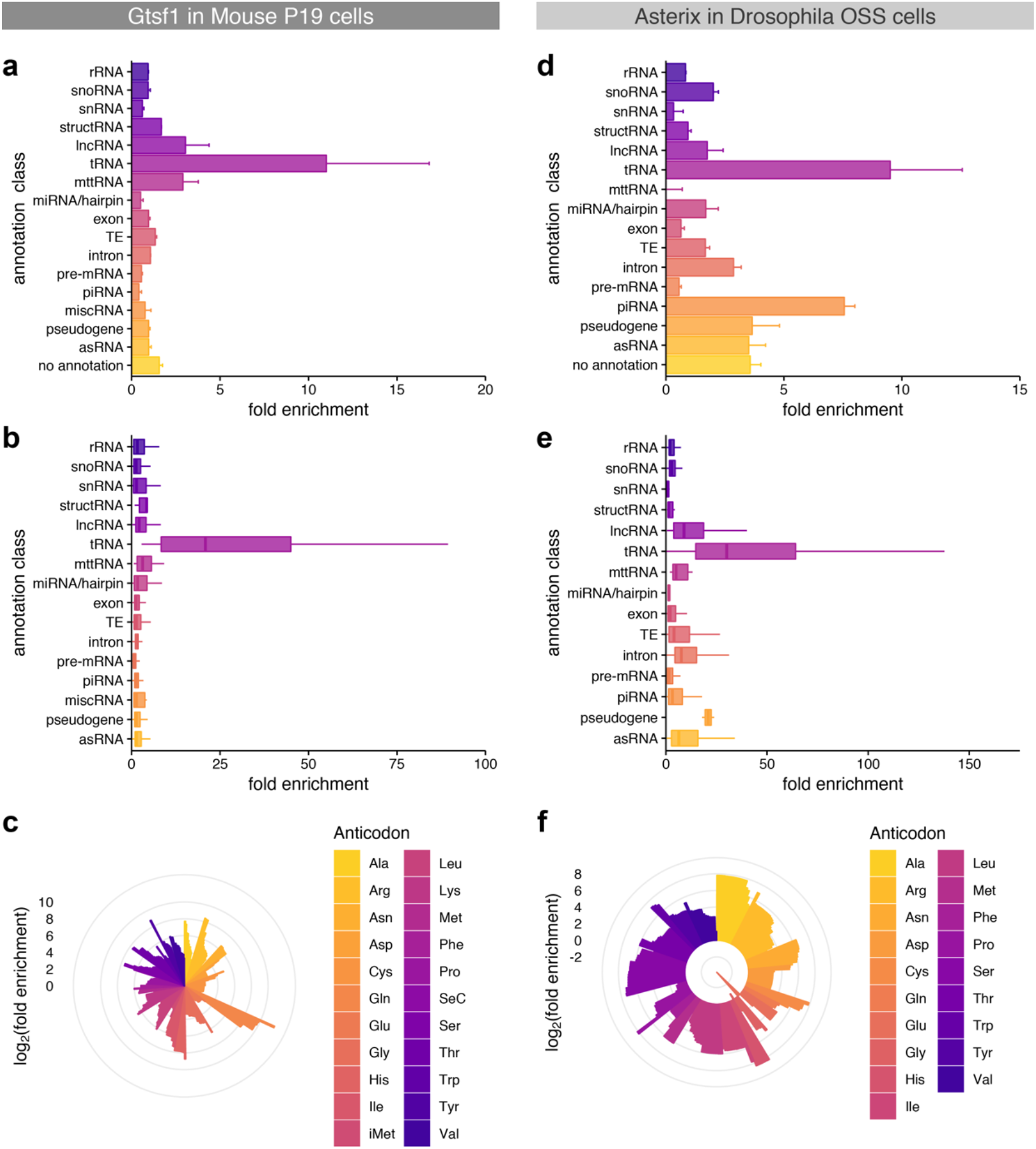
Asterix/Gtsf1 specifically and directly binds tRNAs in cellular contexts. **(a, d) Gene class enrichment of Asterix/Gtsf1-bound RNAs.** The fold enrichment of each annotation class in eCLIP experiments for (a) mouse Gtsf1 in P19 cells and (d) *Drosophila* Asterix in OSS cells is shown as a bar chart. Values indicate the average fold enrichment for two replicate libraries. Error bars indicate the standard error. **(b, e) Annotation enrichment distribution plots.** Fold enrichment distributions among gene annotations within each class are displayed as box plots for (b) P19 mouse and (e) *Drosophila* OSS eCLIP experiments. **(c, f) Fold enrichment scores per tRNA, sorted by anticodon.** tRNA enrichment for (c) P19 mouse and (f) *Drosophila* OSS eCLIP experiments plotted as log_2_(fold enrichment) on the radial bar graph. Multiple bars of the same colours indicate distinct gene annotations for that anti-codon.

We repeated the informatic analysis to test the robustness of these results (Supplemental Fig. 5). We disfavoured tRNA enrichment by setting tRNAs as lowest priority in our annotation ranking (Supplemental Fig. 5b). Additionally, we developed a complementary informatic pipeline that built an alternative gene model to accommodate multimapping reads and then used tools such as DESeq2^32^ to quantify enrichment (Supplemental Fig. 5c). While the absolute strength of the enrichment signals did vary among these analyses, each of these workflows supported the conclusion that tRNAs were the most enriched annotation class.

### Asterix directly binds tRNAs in a *Drosophila* OSS cells

To date, the most productive model organism for dissecting piRNA biology has been *Drosophila melanogaster*. This is due not only to well-established genetic tools but also to the availability of an ovary-derived cell culture line (ovarian somatic sheath cells; OSS) with an intact primary piRNA silencing pathway. The requirement of Asterix for effective transposon silencing in *Drosophila*—and more specifically this cell line—is well-established as the role of this protein in piRNA biology was discovered in OSS cells^9–11^.

Therefore, to compare our observations from mouse Gtsf1 to *Drosophila* Asterix and establish a framework for better cross-referencing observations between mammals and flies, we similarly performed eCLIP experiments in OSS using transfected, Strep-tagged Asterix. Once more, tRNAs were found to be highly enriched both as a class and individually (Fig. 2c-e).

Finally, to verify that these findings were not due to over-expression artefacts, we performed eCLIP experiments using FLAG-tagged Asterix under the control of its endogenous promoter. Again, tRNAs were enriched both individually and as a class (Supplemental Fig. 6). Interestingly, some piRNA enrichment was also observed in this experiment, which may be explained by Asterix’s known association with piRISC complexes^11,17,33^, however, unlike tRNAs his was not found as universally across piRNA annotations.

### Gtsf1 binds tRNAs in the D-arm

To gain insight into the interaction between tRNA and Gtsf1, we further scrutinized the eCLIP data. In eCLIP, a pileup of read ends is expected at the cross-linking site, presumably due to interference from the crosslink with reverse transcription during preparation of the library. Analysis of library 5′ ends can thus be used to inform potential sequence motifs that are specifically engaged with the crosslinked protein.

An initial analysis of genomic sequences in the vicinity of library 5′ ends did not reveal obvious binding motifs. With the apparent preference for tRNAs as a Gtsf1 ligand, and recognizing that tRNAs are highly structured, we hypothesized that RNA-binding by Gtsf1 could be driven by structural determinants perhaps more so than by RNA sequence.

To test this, the 5′-end positions of mapped tRNA reads were plotted as a histogram on a model tRNA 73 nucleotides in length (not including the CCA-tail) and scored according fold enrichment weighted by Z-score (Fig. 3a). Using the analysis which retained the most tRNA reads, we were able to identify two high-scoring sites at nucleotides 18 and 22 in the D-arm (Fig. 3b).

**Figure 3.**
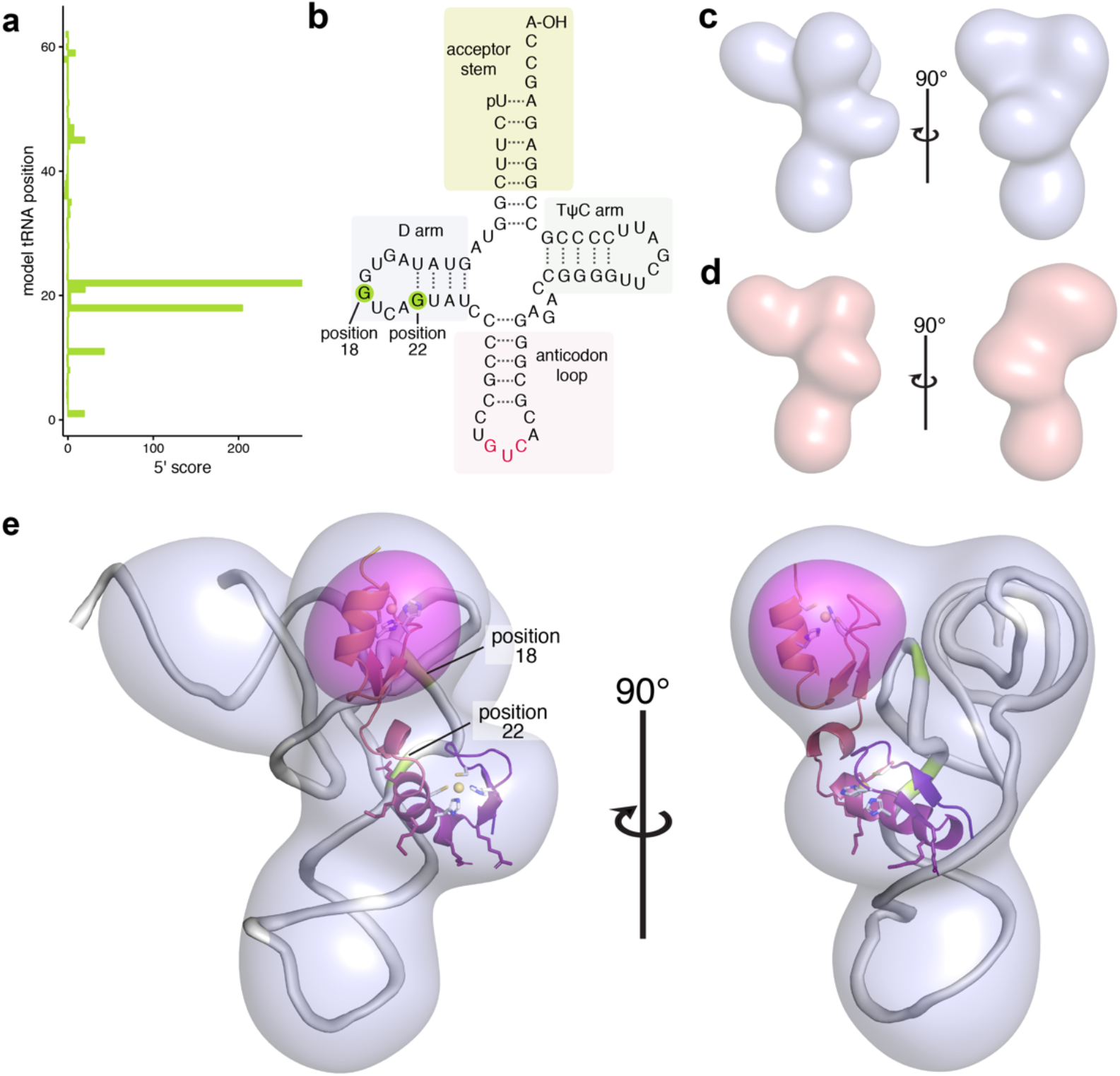
Structure of mouse Gtsf1 in complex with tRNA. (a,b) Mapping of favoured RNA crosslinking sites. (a) Analysis of tRNA reads indicates two preferred cross-linking sites that correspond to nucleotides 18 and 22 of a model tRNA. (b) These sites are in the tRNA D arm and are highlighted (green) on a tRNA secondary structure diagram. **(c,d) Cryo-EM reconstructions of mouse Gtsf1 bound to co-purifying tRNA.** (c) Reconstruction of full-length MmGtsf1 in the presence of co-purifying RNA. The reconstruction is filtered to 10 Å and displayed with a 3.2σ cutoff. (d) Reconstruction of a truncation construct of MmGtsf1 (only the first zinc finger; residues 1-45) in the presence of co-purifying RNA. Filtered to 10 Å and displayed with a 1.4σ cutoff, this reconstruction shows a very similar shape to that observed in (c), but lacking one lobe of the structure. **(e) Difference map and structure.** The full-length reconstruction (light blue) is shown with modelled tRNA (grey), the Gtsf1 NMR structure (core residues 13-72; coloured as in Fig. 1c), and the difference map calculated from reconstructions (c) and (d) (pink surface; contoured at 5.8σ). Positions of favoured tRNA crosslinking are highlighted in green and labelled. Residues of ZnF1 which are important for binding are shown as sticks, coloured according to their position in the ribbon diagram. This alignment accommodates both molecules within the full-length reconstruction, positions the RNA-binding residues of MmGtsf1 ZnF1 near to the tRNA, places the most highly crosslinked RNA residues at the protein interaction surface, and identifies the placement of ZnF2.

### Structure of Gtsf1 in complex with tRNA

Having characterized both Gtsf1 and the RNAs that it interacts with in several contexts, we next aimed to determine a structure of the protein-RNA complex. Initial NMR experiments on Gtsf1 reconstituted with RNA ligands showed evidence of binding, but were hampered by poor quality spectra which likely resulted from decreased molecular tumbling. Attempts at crystallization met with similar difficulties, presumably due to the inherent flexibility present in the protein structure. Although the molecular weight for the complex is only ~45 kDa (19 kDa for Gtsf1 and 25 kDa for a typical tRNA), we speculated that the electronic density of the bound RNA could allow for structure determination using cryo-electron microscopy (cryo-EM).

We subjected recombinantly expressed mouse Gtsf1 from *Sf9* cells, which as mentioned co-purifies with endogenous RNA, to cryo-EM. Given the relatively small size of this complex, the presence of disordered regions in the protein, and the fact that the sample included a heterogeneous population of RNAs, we opted to image this material at 200 keV (rather than the customary 300 keV) to increase contrast and aid in particle picking. Nearly 5,000 micrographs were collected and resulted in almost 500,000 particles (see Methods).

From these data, we were able to obtain a low-resolution reconstruction (Fig. 3c, Supplemental Fig. 7a-b) which had the dimensions and shape of a tRNA with two additional domains. To more accurately orient the Gtsf1 structure into the reconstruction, we applied a similar workflow to an even smaller complex, comprising only ZnF1 in complex with co-purifying RNAs (estimated total molecular weight of 31 kDa) (Fig. 3d, Supplemental Fig. 7c). By comparing the two reconstructions, we were able to generate a difference map to deduce the location of ZnF2 (Fig. 3e, fuchsia surface).

Ultimately, we were able to place tRNA and the zinc finger domains of mouse Gtsf1 into the cryo-EM map, with little residual density, resulting in a low-resolution structure of the complex. Consistent with the biochemical analysis, ZnF1, the domain primarily responsible for binding RNA, formed an interface with most probable crosslinking tRNA nucleotides. The second zinc finger extended toward the tRNA acceptor stem.

### Asterix knockout predominantly affects LTR-class transposons

To understand how binding of Asterix/Gtsf1 to tRNA might be involved in piRNA silencing and repression of transposon expression, we noted that certain groups of retroviruses and retrotransposons require host tRNAs as primers for their replication by reverse transcription^34,35^. Such retrotransposons belong to the LTR (long terminal repeat) family and are characterized by the presence of repeated DNA sequences that flank the transposon body.

In order to transpose, LTR transposon transcripts must be reverse-transcribed. The reverse transcriptase enzyme requires priming, which is most often accomplished using host tRNAs recognizing a primer binding site (PBS) immediately downstream of the 5’ LTR. This dependence of host tRNA recognition thus makes the PBS a conspicuous feature of LTR transposons, which can indeed be exploited for LTR recognition, as has been shown with tRNA fragments^35,36^.

We reasoned that if Asterix/Gtsf1 is indeed using tRNAs to recognize LTR transposon transcripts, then this class of transposons should be highly affected by Asterix/Gtsf1 knockdown. Re-analysis of RNA sequencing data from Asterix knockout flies^11^ supported this finding and indicated that both in absolute read counts and in distributions of fold changes among loci, LTR retrotransposons were indeed the most affected transposon class (Fig. 4a-b).

**Figure 4.**
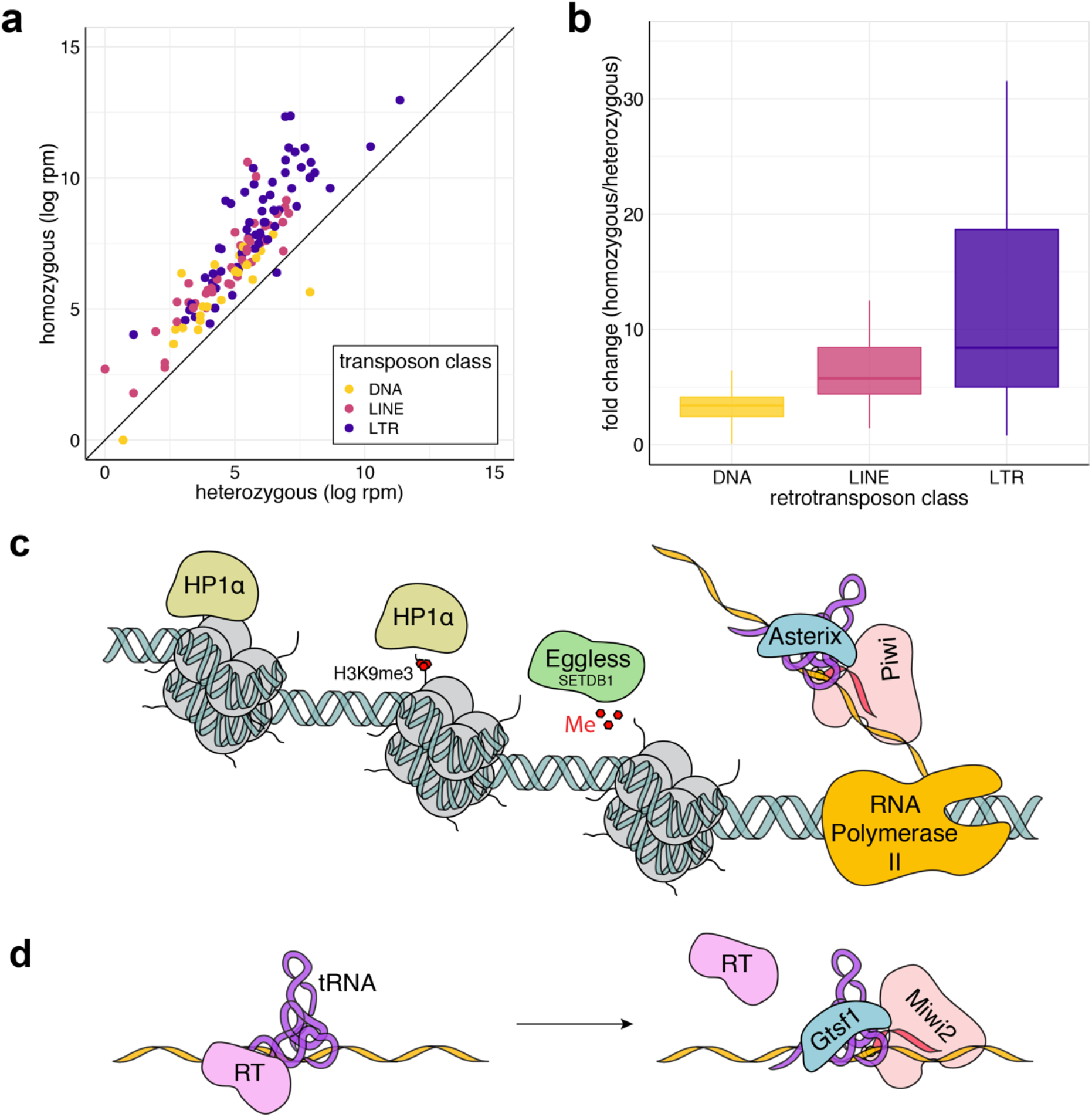
LTR transposons (a class that relies on tRNAs for retrotransposition), are preferentially de-repressed upon Asterix knockout. **(a) Comparison of transposon levels in Asterix knockout versus Asterix heterozygous flies.** Transposon expression as determined by RNA-seq in mutant versus heterozygous flies is plotted, colour-coded by transposon class. **(b) Transposon fold change distribution plots.** Fold changes among gene annotations within each transposon class from are displayed as box plots. **(c,d) Models for the role of Asterix/Gtsf1 in silencing.** (c) In the nucleus, Asterix may utilize tRNAs to recognize primer binding sites in nascent transposon transcripts. Enhanced recruitment of piRISCs (pink) could then be achieved through interactions with Asterix’s C-terminal tail. This leads to further recruitment of histone modification enzymes (such as Eggless/SETDB1; green) and eventual binding of HP1α (beige) to establish heterochromatin. (d) In the cytoplasm, similar interactions between Gtsf1, tRNAs, transposons, and Piwi proteins may also occur to displace reverse transcriptase and/or enhance the slicing activity of cytoplasmic piRISCs.

## Discussion

Several lines of evidence now point to Asterix/Gtsf1 as a *bone fide* tRNA-binding protein: the presence of CHHC zinc fingers that in other proteins bind structured RNAs, co-purification of tRNAs from the recombinant expression of Gtsf1, the ability to abolish these interactions with individual Gtsf1 point mutants, and direct-binding of Gtsf1 to tRNAs in multiple relevant cell culture systems.

Taken together with the marked effects of Asterix knockout on LTR retrotransposons and the evolutionary history that LTR retrotransposons share with tRNAs, tRNA-binding by Asterix/Gtsf1 suggests that these proteins are co-opting molecular epitopes of tRNAs to facilitate transposon silencing.

In the nucleus, where Asterix/Gtsf1 localizes in both mice and flies, LTR tRNA primer binding could be used to augment the specificity of Piwi/MIWI2 (both of which have identified as binding partners^16^) through piRNA base-pairing interactions with transposons (Fig. 4c). In the cytoplasm, Gtsf1 could act to displace or interfere with tRNA-primer/reverse transcriptase engagement and limit retrotransposon replication, while simultaneously assisting recruitment of Piwi partners for ping-pong processing (Fig. 4d). It remains to be understood however, both in typical retroelement replication and in its inhibition through this proposed mechanism, how tRNA unwinding is accomplished, whether it be by additional co-factors or simply part of the dynamic nature the acceptor stem^37^. In both of these cases, initial direction of Asterix/Gtsf1 to the appropriate cellular compartment could be accomplished by its known association with cognate Piwi proteins^16,19^ after which Asterix/Gtsf1 aids in enhancing target recognition.

As RNA interference pathways are studied across many species and cell types, variations on several themes continue to emerge. In addition to the most obvious presence of a small-RNA loaded RISC as the central component of the pathway, complexes that establish multivalent interactions with silencing targets are also prevalent—GW182-mediated recruitment of Ago2 in humans, and the RITS complex in *S. pombe* are prime examples^38–40^ (Supplemental Fig. 8). Moreover, GTSF-1 in *C. elegans* has been demonstrated as critical in the formation of a functional RNA-dependent RNA Polymerase Complex (RDRP) where it is believed to aid in the assembly of RNA silencing complexes.^41^ These multipartite binding platforms confer enhanced molecular specificity while also allowing flexibility in the repertoire of silencing targets. In this case, our findings suggest that Gtsf1/Asterix exploits a key vulnerability in many retro elements: their dependence on host tRNAs for their replication.

## Methods

### Cloning

#### Overview

In order to screen for well-behaved targets for recombinant protein expression, a panel of constructs was generated from *H. sapiens, M. musculus*, and *D. melanogaster* Gtsf1 cDNAs (codon-optimized for expression in *Sf9*) by SLIC (sequence- and ligation-independent cloning). These constructs presented various N- or C-terminal tags for enhanced expression and purification using either *E. coli* or insect cell culture systems. In addition, natural sequences of the *D. melanogaster* and *M. musculus* proteins were used for transfection in eCLIP experiments. The sequence of each construct was verified by GenScript. Constructs presented in this work are described in further detail below and summarized at the end of this section.

#### Constructs for structure determination by NMR (expression in E. coli)

Among the constructs screened for expression in *E. coli*, a fragment corresponding to the first 115 residues of MmGtsf1 with a C-terminal His_6_-tag showed highest expression and produced sufficiently soluble material for structure determination by NMR. To prevent aggregation over the duration of NMR data collection, four of the cysteines (those not involved in zinc chelation) were mutated to serine. These constructs were cloned into the vector pET-22 and also included TEV-cleavable linker for His_6_-tag removal (MmGtsf1-115-TEV-His and MmGtsf1-115-TEV-His C28S, C76S, C100S, C103S).

#### Constructs for RNA binding studies (expression in Sf9)

Constructs were similarly screened for expression in *Sf9* cells. Data presented for RNA interaction studies include the full-length protein (167 residues), point mutants, and truncations as indicated in each figure. All *Sf9*-derived material included a C-terminal Strep_2_-tag and TEV-cleavable linker and was cloned in to the vector pFL for baculoviral-induced insect cell culture.

#### Constructs for eCLIP

MmGtsf1 cDNA (not codon-optimized) was obtained from GensScript (Accession Number NM_028797.1; Clone ID: OMu06141D) and subcloned by SLIC into the vector pEFα under the control of the EF1α promoter. A C-terminal TEV-Strep_2_ tag was included to allow for affinity pull-down during eCLIP processing. pmaxGFP (Lonza) was used as a transfection control. pEFα plasmid was a kind gift from A. Schorn and R. Martienssen.

### Expression and Purification

#### Expression in E. coli for NMR structure determination

Target constructs were transformed into BL21-CodonPlus (DE3)-RIPL. Isotopically-labelled (U-^15^N and/or U-^13^C) protein was produced using M9 media supplemented with ^15^NH_4_Cl and/or ^13^C-labeled glucose. Cells were grown at 37°C to a culture density of approximately 0.7. Protein expression was induced with 1 mM IPTG (final concentration) and proceeded for 3.5 hours.

#### Purification

Cells were harvested by centrifugation at 4000*g*, resuspended in lysis buffer (50 mM sodium phosphate, pH 8.0, 50 mM NaCl, 10 mM imidazole; ~20 mL per litre culture), and lysed by sonication. The cell lysate was clarified by ultracentrifugation at 125,000*g* for 1 h after which the supernatant applied to a Ni-NTA column equilibrated with lysis buffer. The column was washed (50 mM sodium phosphate, pH 8.0, 300 mM NaCl, 40 mM imidazole) and the protein then eluted (50 mM sodium phosphate, pH 8.0, 300 mM NaCl, 200 mM imidazole). To prevent precipitation and proteolysis, DTT was added to the elution at a final concentration of 10 mM and EDTA at a final concentration of 1 mM. The C-terminal His_6_-tag was then removed by overnight treatment with TEV protease (1:25 mass ratio of protease:target) at 4°C. The cleaved protein was further purified by ion-exchange chromatography (MonoQ column) in a buffer of 25 mM Tris, pH 8.0, and 2 mM DTT with a NaCl gradient from 0 to 1 M. MmGtsf1-115 eluted approximately between 17 and 24 mS. Peak fractions were pooled, concentrated, and used for further purification by gel filtration chromatography (Superdex75 increase) in 50 mM MES, pH 6.5, 200 mM NaCl, and 5 mM TCEP. Peak fractions were pooled, concentrated and mixed with ZnCl_2_ (2:1 molar ratio Zn^2+^:protein) and MgCl_2_ (4:1 molar ratio Mg^2+^:protein). Upon addition of ZnCl_2_, the protein solution became temporarily turbid, but clarified upon gentle mixing. For NMR structure determination, sodium azide was added at a final concentration of 0.02% as a preservative. Typical yields were 2-3 mg of purified protein (>98% pure as assessed by SDS-PAGE) per litre culture.

#### Expression in Sf9 and RNA pull-down

Indicated constructs (each with a C-terminal TEV-Strep_2_ tag) were cloned into the vector pFL for expression in *Sf9* cells using the baculovirus expression system. After expression, cells were harvested by centrifugation at 1000*g*, resuspended in lysis buffer (50 mM Tris, pH 8.0, 100 mM KCl, 1 mM DTT) (~20 mL per litre culture), and lysed by sonication. The cell lysate was then clarified by ultracentrifugation at 125,000*g* for 1 h and the supernatant applied to a *Strep*-Tactin (IBA) column equilibrated with lysis buffer. The bound MmGtsf1 proteins were subsequently washed with lysis buffer, further washed with lysis buffer containing 2 mM ATP, and finally eluted in lysis buffer containing 5 mM D-desthiobiotin. Protein purity was assessed by SDS-PAGE. Co-purifying nucleic acids were isolated by phenol:choloform extraction, precipitated with ethanol, then assessed by Urea-PAGE.

### Characterization of Co-purifying *Sf9* RNAs

#### Initial nucleic acid characterization

After phenol:chloroform extraction and alcohol precipitation, pulled-down nucleic acids were characterized by treatment with RNase, DNase, or by alkaline hydrolysis. For each treatment, approximately 50 ng of nucleic acid was mixed with either RNase A (Ambion; 1 μg), DNase I (Zymo Research; 0.1 units), or 1 μL of 1 M sodium hydroxide in total volume of 40 μL under suitable buffer conditions (10 mM Tris, pH 8.0, 1 mM EDTA for RNase A treatment; no added buffer for alkaline hydrolysis treatment; 10 mM Tris, pH 7.6, 2.5 mM MgCl_2_, 0.5 mM CaCl_2_ for DNase I treatment). Murine RNase inhibitor (NEB; 40 units) was included in all conditions with the exception of the RNase A treatment. Samples were incubated for 15 minutes at 37°C for nuclease treatments or 70°C for alkaline hydrolysis. After treatment, the sodium hydroxide was neutralized by the addition of 1 μL of 1 M hydrochloric acid. As a control, a 50 nucleotide DNA duplex was treated under the same set of conditions. All samples were the denatured and assessed by 12% Urea-PAGE.

#### sRNA library preparation

Affinity co-purifying nucleic acids which bound to MmGtsf1 during expression in *Sf9* were separated from the protein by ion exchange chromatography (Mono Q column, as described above, eluting between 45 and 55 mS). Peak fractions were pooled, and the RNA isolated by phenol:chloroform extraction and alcohol precipitation. Small RNA libraries were prepared using the SMARTer smRNA-Seq Kit for Illumina sequencing (Takara). Size-selection was performed using Blue Pippin 2% agarose gel cassettes (Sage Science). All libraries were assessed by fluorometric quantification (Qubit 3.0) and by Bioanalyzer chip-based capillary electrophoresis. The average fragment size was 228 bp with most insert sizes ranging from 20-100 bp. Libraries were pooled in equimolar ratios according to their quantification (determined above). Single-end reads with two 8-basepair barcodes were generated on an Illumina NextSeq resulting in approximately 10 million reads per library.

#### sRNA library data processing

Owing to the incomplete assembly of the *Sf9* genome and the lack of annotations, processing for sRNA was straightforward, but limited. Reads were first trimmed to remove sequences appended during library preparation (adapters, polyA sequences at the 3′ end, as well as the first three nucleotides after the adapter at the 5’ end). Removal of the polyA sequence was performed using a custom script (*polyA_trim.py*). Reads were then filtered based on size and quality scores. Reads in the processed libraries were collapsed and the most abundant sequences were manually inspected.

### eCLIP Library Generation

#### P19 cell culture

P19 (mouse embryonal carcinoma; ATCC CRL-1825) cells were maintained in minimum essential medium with ribonucleosides and deoxribonucleosides (Gibco), supplemented with bovine calf serum and fetal bovine serum (7.5% and 2.5% final concentration, respectively) (Seradigm). Cells were cultured at 37°C in a 5% CO_2_ atmosphere. Cultures were monitored for *Mycoplasma* contamination using the MycoAlert Mycoplasma Detection Kit (Lonza). *Mycoplasma* contamination was not detected. The identity of the cultured cells was confirmed by short tandem repeat (STR) profiling, serviced by ATCC.

#### OSS cell culture

Drosophila OSS (ovarian somatic sheath) cells were maintained in OSS medium (Shields and Sang M3 Insect Medium [Sigma-Aldrich] supplemented with approximately 5 mM potassium glutamate, 5 mM potassium bicarbonate, 10% heat-inactivated fetal calf serum [Invitrogen], 10% fly extract [Drosophila Genomics Resource Center], 2 mM reduced glutathione [Sigma-Aldrich], 1x GlutaMAX [Gibco], 0.01 mg/mL human insulin [Sigma-Aldrich], and an antibiotic-antimycotic [Gibco] consisting of penicillin, streptomycin, and Amphotericin B). Cells were cultured at ~23°C. Cultures were monitored for *Mycoplasma* contamination using the MycoAlert Mycoplasma Detection Kit (Lonza). *Mycoplasma* contamination was not detected. OSS cells with Asterix C-terminally FLAG-tagged at its native locus (Asterix-GFP-FRT-Precission-V5-FLAG_3_-P2A) were provided by the lab of J. Brennecke and cultured in the same way as unmodified OSS cells.

#### Transfection of MmGtsf1-TEV-Strep into P19 cells

For transfection, P19 cells were grown to 75% confluency in 150 mm culture dishes. Four hours prior to transfection, the medium was refreshed. To transfect, 30 μg of DNA (either MmGtst1-TEV-Strep in pEF or eGFP in pMAX [transfection control]) was premixed with 60 μL of X-tremeGENE HP DNA transfection reagent (Roche) in serum-free medium for 15 minutes. After a 15-minute incubation, this mixture was added to the cultures. Sixteen hours post-transfection, the cells were visibly perturbed and the medium was again refreshed. Expression of eGFP in the transfection control was confirmed by UV microscopy. Forty-eight hours post-transfection, the cells were rinsed with ice-cold phosphate-buffered saline (PBS) and taken for processing.

#### Transfection of Asterix-TEV-Strep into OSS cells

For transfection, OSS cells were grown to 75% confluency in 150 mm culture dishes. Four hours prior to transfection, the medium was refreshed. To transfect, 50 μg of DNA (either Asterix-TEV-Strep in pAWG or pAGW [transfection control]) was premixed with 15 μL of Xfect Polymer transfection reagent (Takara) in 500 μL Xfect buffer. OSS medium was removed from the cells and replaced with Shields and Sang M3 Insect Medium supplemented only with potassium bicarbonate and potassium glutamate. After a 10-minute incubation of the DNA with the transfection reagent, the transfection mixture was added to the cultures. Two hours post-transfection, the M3 medium was removed and replaced with fully-supplemented OSS medium. Expression of GFP in the transfection control was confirmed by UV microscopy. Seventy-two hours post-transfection, the cells we rinsed with ice-cold phosphate-buffered saline (PBS) and taken for processing.

#### Library preparation

eCLIP Libraries were prepared essentially as in Van Nostrand *et al.*^25^ with the following parameters and modifications. UV cross-linking was performed at 254 nm for ~45 seconds (400 mJ) in an HL-2000 Hybrilinker. For MmGtsf1-TEV-Strep in P19 cells and Asterix-TEV-Strep in OSS cells, protein pull-down was accomplished using MagStrep “type 3” XT beads (IBA) with 50 μL of bead resuspension used per sample. Asterix-GFP-FRT-Precission-V5-FLAG_3_-P2A, pull-down was similarly accomplished with Anti-FLAG M2 magnetic beads (Sigma-Aldrich).

Final libraries were amplified and barcoded using Illumina compatible primers as described below.

|  |  |
| --- | --- |
| Non-crosslinked input | D504, D701 |
| Crosslinked input (replicate 1) | D504, D702 |
| Crosslinked input (replicate 2) | D501, D703 |
| Non-crosslinked IP (background) | D503, D701 |
| Crosslinked IP (replicate 1) | D502, D703 |
| Crosslinked IP (replicate 2) | D503, D704 |

For samples from mouse P19 cells, 8 amplification cycles were used for the inputs and 14 cycles for the IPs. For samples from *Drosophila* OSS cells, 13 amplification cycles were used for the inputs and 18 cycles for the IPs.

For quality control, all libraries were assessed by fluorometric quantification (Qubit 4.0) and by Bioanalyzer chip-based capillary electrophoresis. The average fragment size was typically 240-250 bp with most insert sizes ranging from 15-200 bp. A detailed version of the complete eCLIP library preparation is available upon request.

#### Next-generation sequencing

Libraries were pooled in equimolar ratios according to their quantification (determined above). Paired-end reads with two, 8-basepair barcodes were generated on an Illumina NextSeq resulting in approximately 100 million paired-end reads (~15-20 million reads per library).

### eCLIP Processing

#### Rationale

Based on our previous findings when sequencing endogenous *Sf9* RNAs copurifying with recombinantly-expressed MmGtsf1, we surmised that it would be necessary to include multi-mapping reads in our analysis pipeline. This stems from the fact that many of the RNA species of interest arise from known multi-mapping regions (tRNAs, transposable elements, and piRNA clusters).

#### Summary

The pipeline begins with demultiplexed paired-end libraries. Given that most all of the paired-end reads were short enough to overlap, they were joined into single sequences using FLASH^42^. Sequencing adapters were then trimmed, PCR duplicates removed, and the reverse complement of the read (corresponding to the sense strand of the original RNA) was taken for downstream processing. Identical reads were collapsed and counted, then mapped to the genome using STAR^26^. The aligned reads were then annotated and filtered based on feature type. Full descriptions of custom scripts accompany the code on GitHub (see Data Availability).

### Gene Annotations

As many of the gene classes of interest have dedicated communities of their own (tRNAs, miRNAs, piRNAs, and transposons), we incorporated these multiple annotation sources into the workflow. The sources of annotations are listed below for both the mouse and *Drosophila* analyses. In brief, annotations from each source were compared, matched when possible, and if matched the outer bounds of each annotation were taken. The resulting composite annotations have been deposited on GitHub (see Data Availability).

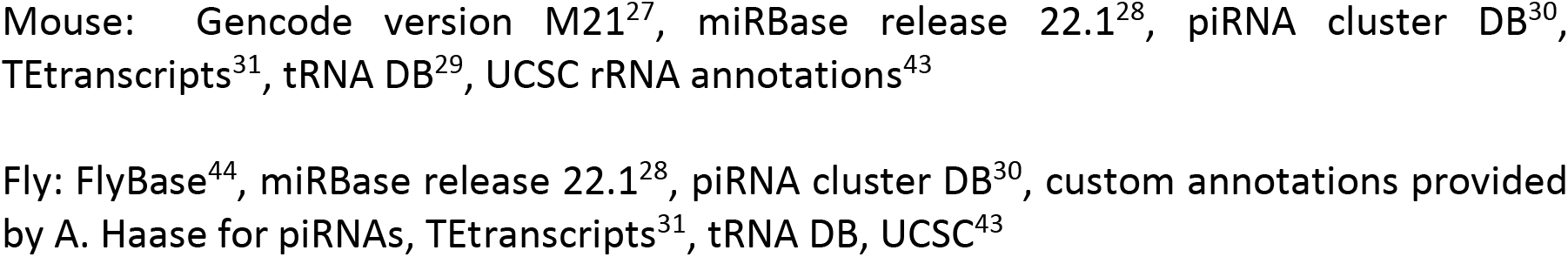

### tRNA Analysis

Following multi-mapping normalization, reads belonging to the tRNA annotation class were further characterized. To begin, the size of each tRNA annotation was scaled to a “model tRNA” size of 73 nucleotides. Each tRNA read was then re-mapped to its annotation, now scaled to the model tRNA length. By aggregating all tRNA-mapping reads, we were able to generate histograms of read statistics (5′ end, 3′ end, read length, and nucleotides covered). It is expected that eCLIP reads will have a pileup at their 5′ end corresponding to the cross-linking site. We scored this pileup by determining the fold enrichment for each metric (essentially calculated as [IP – background] / input) and weighting it by its Z-score.

### NMR Spectroscopy

#### Instrumentation

NMR spectroscopy was performed using Bruker AVANCE500 (New York Structural Biology Center, NYSBC), DRX600 (Columbia University), AVANCE700 (NYSBC), AVANCE800 (NYSBC), and AVANCE900 (NYSBC) NMR spectrometers equipped with 5 mm cryoprobes.

#### Sample preparation

MmGtsf1 samples were prepared in 50 mM MES, pH 6.5, 200 mM NaCl, 5 mM TCEP, and 2:1 stoichiometric ZnCl_2_, 4:1 stoichiometric MgCl_2_, and 0.02% azide. For data acquisition, samples were either supplemented with a final concentration of 10% D_2_O or lyophilized and resuspended in 99% D_2_O. Sample concentrations were 0.5 mM for the [U-^15^N]-labelled protein and 0.8 mM for the [U-^13^C, U-^15^N]-labelled protein. The sample temperature was calibrated to 298 K using 98% ^2^H_4_-methanol^45^. 100 μM DSS was included in samples for internal referencing of ^1^H chemical shifts, followed by indirect referencing for ^13^C and ^15^N chemical shifts^46^.

#### Resonance assignments

Backbone resonance assignments were obtained using ^1^H-^15^N HSQC, ^1^H-^13^C HSQC, HNCA, HN(CO)CA, HNCO, HN(CA)CO, HNCACB, and HN(COCA)CB experiments^46^. Side chain resonance assignments were obtained using HCCH-TOCSY, HBHA(CO)NH, H(CCO)NH, and (H)C(CO)NH experiments^46^. Spectra were processed using nmrPipe^47^ and analysed using Sparky^48^.

#### Distance restraints

Distance restraints for structure determination were obtained from ^1^H-^15^N NOESY-HSQC, ^1^H-^13^C NOESY-HSQC, and ^1^H-^13^C NOESY-HSQC (with spectral parameters optimized for detection of aromatic spins)^46^. ^1^H-^13^C NOESY experiments were performed for samples prepared in 99% D_2_O.

#### Zinc coordination

Protonation states of histidine residues were determined by long-range HMQC experiments together with the empirical correlation between the chemical shift difference ^13^C_∊1_ - ^13^C_δ2_^49^. H23 and H57 are designated with N_δ1_ coordination, and H33 and H67 are designated with N_∊2_ coordination to the Zn^2+^ ion.

#### Relaxation parameter determination

Backbone ^15^N *R*_1_ relaxation rate constants, ^15^N *R*_2_ relaxation rate constants, and the steady-state {^1^H}-^15^N NOE were measured at 500 MHz (NYSBC) using the pulse sequences of Lakomek *et al.*^50^. *R*_1_ measurements used relaxation delays of 24 (×2), 176, 336 (×2), 496, 656, 816, 976, and 1200 ms. *R*_2_ measurements used relaxation delays of 16.3 (×2) 32.6, 49.0 (×2), 65.3, 97.9, 130.6, 163.2, and 195.8 ms. NOE measurements used a recycle delay of 7 s for the control experiment and 2 s of recovery followed by 3 s of saturation for the saturated experiment. Duplicate relaxation delays were used for error estimation for measurement of ^15^N R_1_ and *R*_2_ relaxation rate constants. Duplicate experiments were used for error estimation for the steady-state {^1^H}-^15^N NOE experiment.

#### Structure determination

Automatic NOESY cross-peak assignments and structure calculations were performed with ARIA 2.3 (Ambiguous Restraints for Iterative Assignment)^51^ using an eight step iteration scheme supported by partial manual assignments of aliphatic/aromatic ^13^C-edited NOESY-HSQC and amide ^15^N-edited NOESY-HSQC spectra, respectively. Less than 10% of all assignments were labelled ambiguous after initial and final ARIA structure calculations. The unambiguous distance restraints output from the automation run was recalibrated by increasing all the upper distance limits by ~10% and further elimination of lone and consistent NOE violations by manually inspecting the lower quality peak assignments. Dihedral angle restraints for residues in the structured zinc finger domains were derived from the analysis of the backbone chemical shifts in TALOS^52^. Structure calculations were performed in two stages by initially excluding Zn^2+^ during automated NOESY cross-peak assignments followed by water refinement of the Zn^2+^-bound structures. The tetrahedral Zn^2+^ metal ion coordination was implemented in CNS 1.1 by adding a CHHC patch with the experimentally verified tautomeric states for the two histidine side chains in the topallhdg5.3.pro file^21,53^. Bond lengths and angles used to define the Zn^2+^-bound CHHC motif in the parallhdg5.3.pro file was obtained from the published structure (PDB 2VY4) of the homologous ribonuclear protein U11-K48^21^. The final ensemble of 20 representative Zn^2+^-bound structures was generated by calculating 500 structures with water refinement in CNS 1.1^54^. Supplemental Table 1 summarizes the final restraints used in the calculations, NMR ensemble statistics, and the overall quality of the structures determined by MolProbity^24^.

#### Local variability analysis

A sliding window of ±3 amino acids was used to align the 20 lowest energy structures to one another in all combinations at each residue. Average RMSDs were calculated for each window’s alignment, then mapped onto the central residue in the window. Residues near the termini included as many residues as possible while maintaining up to 3 residues on either side of the queried residue (e.g. the score for residue 1 derived from RMSDs using residues 1-4 for alignment; the score for the final residue, 115 derived from RMSDs using residues 112-115).

### Cryo-electron Microscopy

#### Sample preparation

Affinity-purified MmGtsf1-TEV-Strep constructs from *Sf9* (which included co-purified RNAs) at ~0.25 mg/mL in elution buffer (50 mM Tris, pH 8.0, 100 mM KCl, 1 mM DTT, 5 mM d-desthiobiotin) were first cross-linked at 254 nm for ~45 seconds (400 mJ) in an HL-2000 Hybrilinker. It should be noted however, that assessment of RNAs by Urea-PAGE following this treatment did not seem to result in significant covalent cross-linking. For cryo-EM grid preparation, 4 μL of solution was applied to a glow-discharged Lacey carbon grid, incubated for 10 seconds at 25°C and 95% humidity, blotted for 2.5 seconds, then plunged into liquid ethane using an Automatic Plunge Freezer EM GP2 (Leica).

#### Data acquisition

Data were acquired on Titan Krios transmission electron microscope (ThermoFisher) operating at 200 keV. Dose-fractionated movies were collected using a K2 Summit direct electron detector (Gatan) operating in electron counting mode. In total, 32 frames were collected over a 4 second exposure. The exposure rate was 7.6 e^−^/pixel/second (approximately 19 e^−^/Å^2^/second), which resulted in a cumulative exposure of approximately 76 e^−^/Å^2^. EPU data collection software (ThermoFisher) was used to collect micrographs at a nominal magnification of 215,000x (0.6262 Å/pixel) and defocus range of −1.0 to −3.0 μm. For the full-length protein construct sample (MmGtsf1-TEV-Strep with RNA), 4,849 micrographs were collected. For the construct containing only the first zinc finger (MmGtsf1-[1,45]-TEV-Strep with RNA), 2,461 micrographs were collected.

#### Micrograph processing and 3D reconstruction

Real-time image processing (motion correction, CTF estimation, and particle picking) was performed concurrently with data collection using WARP^55^. Automated particle picking was initiated with the BoxNet pretrained deep convolutional neural network bundle included with WARP that implemented in TensorFlow. Following this first round of particle picking, the particle selections on ~20 micrographs were manually inspected and adjusted. This process was iterated one additional time. For the full-length construct, a particle diameter of 100 Å and a threshold score of 0.6 yielded 495,299 particle coordinates for the full-length construct. These particles were then subjected to a 2D classification in cisTEM^56^ after which a subset of 346,643 particles were used for *ab initio* reconstruction and autorefinement in cisTEM. For the truncation construct, a particle diameter of 100 Å and a threshold score of 0.5 yielded 159,646 particle coordinates. These were then taken for 2D classification in cisTEM^56^ after which a subset of 96,036 particles were used for *ab initio* reconstruction and autorefinement. After refinement, structures of tRNA— modelled incorporating the sequence of the most abundantly pulled-down RNA from *Sf9* expression of MmGtsf1-TEV-Strep (Supplemental Table 2) and generated with RNAComposer^57,58^—and the zinc finger domains of MmGtsf1 were manually placed in the reconstructed volume based on the molecular shapes and the likely interaction surfaces as defined by mutagenesis data and most probable eCLIP cross-linking sites.

#### Difference map calculation

Reconstructed volumes for the full-length and truncated MmGtsf1 constructs (both with co-purifying RNA as described above) were filtered to 10 Å with cisTEM^56^. Using SPIDER^59^, a 90 pixel (~56 Å) radius mask was applied to the filtered volumes after which each was normalized and aligned. This map for the truncation construct was then subtracted from the corresponding full-length map (MmGtsf1-TEV-Strep with RNA).

### Figures

Figures of molecular models were generated using PyMOL^60^. Electrostatic surface calculations were performed with APBS^61^ with a solvent ion concentration of 0.15 M using the AMBER force field. Superpositioning of structural homologs was performed by the DALI server^62^. Conservation analysis was performed using the Consurf server^63^. Co-evolution analysis was performed using the Gremlin server^23^. Graphs were produced in R^64^ using the ggplot2 package^65^.

## Data Availability

Coordinates and NMR data have been deposited in the Protein Data Bank (PDB: 6X46) and the Biological Magnetic Resonance Bank (BMRB: 30754). Cryo-electron microscopy maps for complexes isolated from full-length MmGtsf1 protein and ZnF1 domain pull-downs have been deposited in the EMDB (EMD-22040 and EMD-22041, respectively). Sequencing data have been deposited in Gene Expression Omnibus (GEO) repository with accession numbers GSE151110 (Sf9 RNA pull-down), GSE151108 (eCLIP data from P19 cells), GSE151107 (eCLIP data from OSS cells), GSE151109 (eCLIP data from OSS cells using CRISPR-tagged Asterix). Custom gene annotation files and data processing scripts are available on GitHub (https://github.com/jonipsaro/asterix_gtsf1). Intermediate files used for generating gene annotations or processing of the data are available upon request.

## Acknowledgements

We thank Sonam Bhatia, Sara Ballouz, Sarah Diermeier, Paloma Guzzardo, Matt Jaremko, Justin Kinney, Molly Gale Hammell, Greg Hannon, Katie Meze, Felix Muerdter, Kathryn O’Neill, Nikolay Rozhkov, David Spector, and Dennis Thomas for advice. Astrid Haase provided technical advice in culturing OSS cells and custom piRNA cluster gene annotations for *D. melanogaster*. CRISPR-tagged Asterix OSS cells were gifted by Julius Brennecke. P19 cells and advice were provided by Andrea Schorn and Rob Martienssen. We thank Amanda Epstein and visiting students Michael Jacobs, Edward Twomey, and Dexter Adams for technical assistance. Next-generation sequencing was conducted at the CSHL Woodbury Genome Research Center. We thank Life Science Editors for editorial assistance. This work was supported by National Institutes of Health (NIH) grants F32GM097888 (J.J.I.), R01GM050291 (A.G.P.), R35GM130398 (A.G.P.) and T32 GM008281-28 (P.A.O). The 600 and 800 MHz NMR spectrometers were supported by NIH grants S10RR026540 and S10OD016432, respectively. Some of the work presented here was conducted at the Center on Macromolecular Dynamics by NMR Spectroscopy located at the New York Structural Biology Center, supported by NIH grant P41 GM118302. L.J. is an Investigator of the Howard Hughes Medical Institute.

## Author Contributions

J.J.I. conceptualized and performed biochemical, next-generation sequencing, informatic analysis, NMR sample preparation, and cryo-electron microscopy. P.A.O. and S.B. collected and processed NMR data. A.G.P. conceptualized and supervised NMR data collection and assisted with analysis. L.J. conceptualized and supervised biochemical, sequencing and informatic, and cryo-electron microscopy experiments. All authors contributed to manuscript preparation.

## Supplemental Information

Supplementary Information is available for this paper:

Supplemental Figures 1-8

Supplemental Tables 1-2

## Correspondence Statement

Correspondence and requests for materials should be addressed to.

---

**Supplemental Figure 1.**
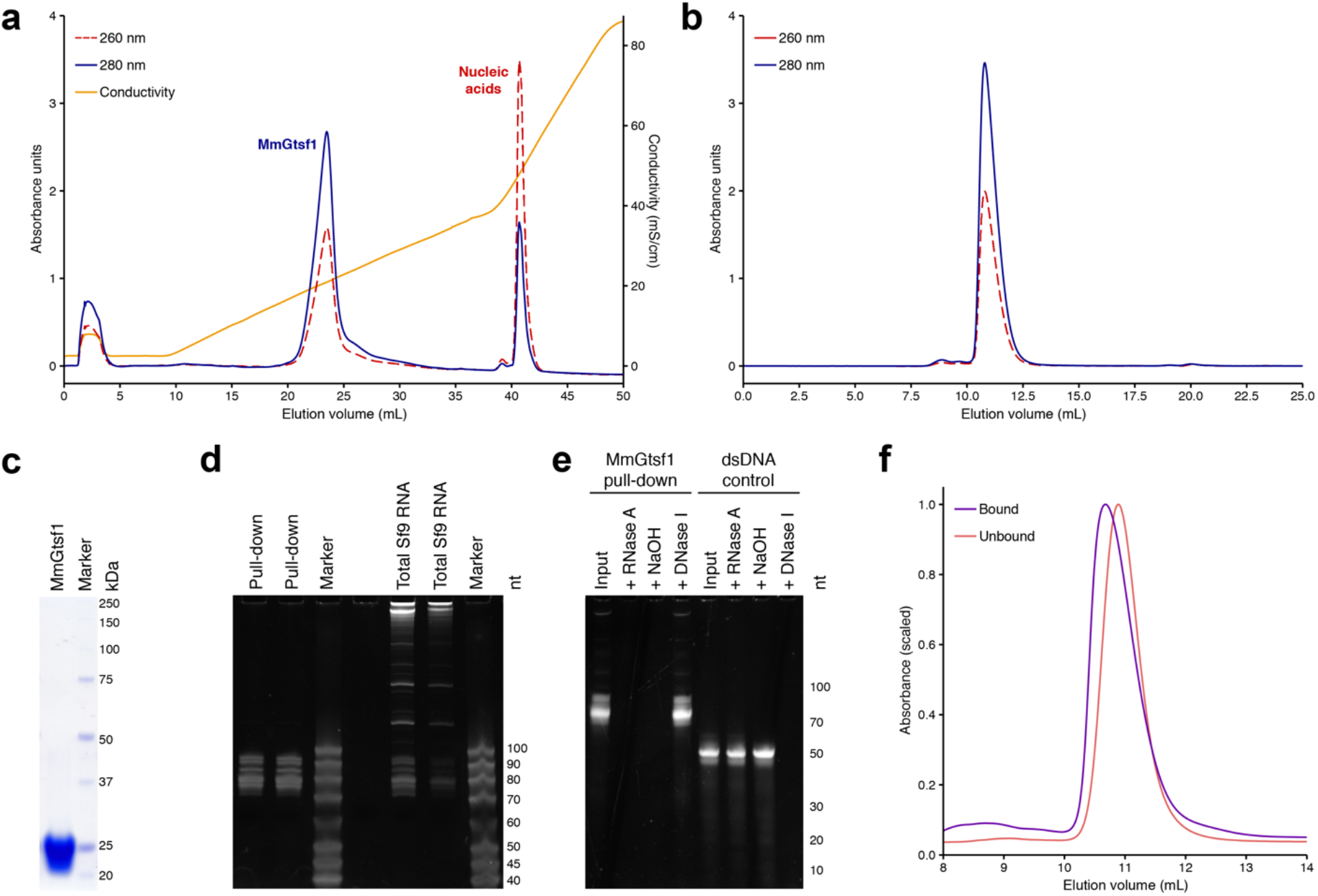
Purification and characterization of MmGtsf1 and co-purifying nucleic acids. **(a) Ion-exchange purification of MmGtsf1 expressed in***Sf9*. The peak eluting at ~25 mS/cm corresponded to MmGtsf1 protein based on the A_λ=260_/A_λ=280_ ratio and verified by SDS-PAGE. The peak eluting at ~50 mS/cm corresponded to co-purifying nucleic acids. **(b) Gel filtration purification of MmGtsf1.** Peak fractions from the ion exchange purification that corresponded to protein (21-24 mL) could be further purified by gel filtration chromatography, eluting as a single peak. **(c) SDS-PAGE assessment of purified wildtype MmGtsf1 expressed in** *Sf9*. SDS-PAGE indicated highly purified material. Approximately five micrograms were loaded. **(d) Urea-PAGE assessment of co-purifying nucleic acids.** Peak fractions from ~40-43 mL contained nucleic acids 70-90 nucleotides in size. Extracted total RNAs from *Sf9* display a much larger size distribution. Two-hundred nanograms of nucleic acid were loaded per lane. **(e) Biochemical determination of nucleic acid identity.** The co-purifying nucleic acids were subjected to various enzymatic and chemical treatments. Both RNase A and sodium hydroxide treatment resulted in degradation of the input material while treatment with DNase I did not, indicating that the co-purifying nucleic acids are RNA (lanes 1-4). Similar treatments of a double-stranded DNA oligonucleotide (lanes 5-8) are included as treatment controls. **(f) Comparison of gel filtration elution profiles.** MmGtsf1 applied to gel filtration chromatography before (purple) and after (pink) removal of co-purifying nucleic acids suggests that a stable complex is originally formed that can be disrupted by ion-exchange purification.

**Supplemental Figure 2.**
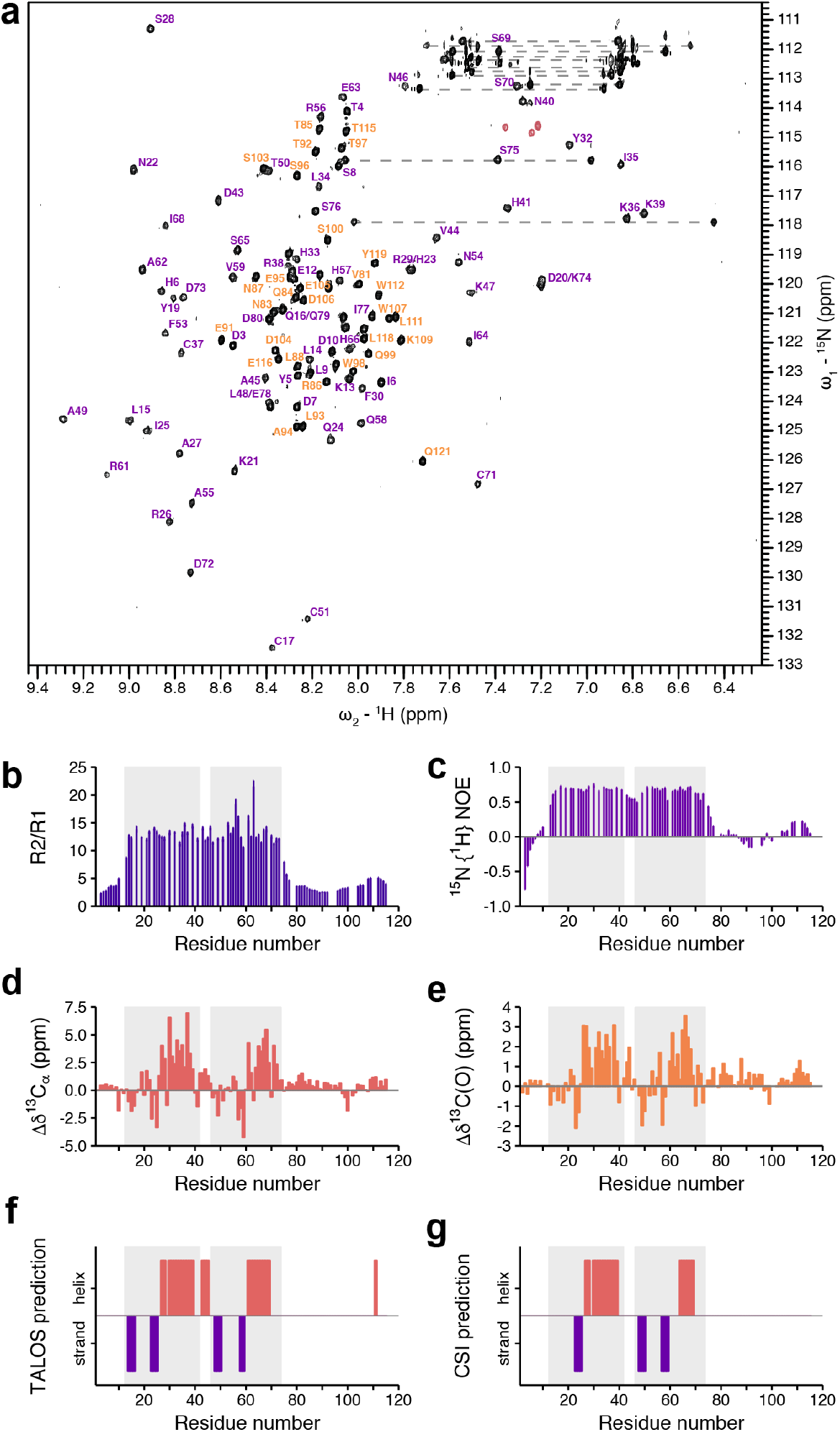
Summarized NMR analysis. **(a) ^**1**^**H-**^s**15**^N HSQC spectrum of MmGtsf1-115.** Backbone amide resonances are labelled using one letter amino acid codes and residue number. Labels for residues for which the structure was determined (1-80) are coloured in purple. Labels for additional residues in the construct (81-115 plus six residues in the C-terminal tag) are coloured orange. Side chain NH_2_ resonances are connected by horizontal lines and are unlabelled. Arginine ^1^H_∊_-^15^N_∊_ resonances are folded in the spectrum. **(b, c) ^15^N spin relaxation assessment of MmGtsf1-115**. Generally uniform, large values of (b) R_2_/R_1_ and (c) the {^1^H}-^15^N NOE for the zinc finger domains indicate that the protein backbone is highly ordered in these regions. Reduced values of R_2_/R_1_ and {^1^H}-^15^N NOE values of the near zero or negative indicate substantial disorder in the N- and C-terminal regions flanking the zinc fingers. The linker between the two zinc fingers also have relaxation data compatible with an ordered backbone. Elevated values of R2/R1 for residues R56 and E63 may indicate a contribution from chemical exchange dynamics. **(d, e) Secondary chemical shifts for MmGtsf1-115.** The secondary chemical shifts μδ, defined as the measured chemical shift minus the residue-specific random coil chemical shift, are shown for (d) ^13^Cα and ^13^C(O) spins. Large positive and negative secondary shifts are indicative of helical or strand (extended) backbone conformations, respectively. Thus, helical elements are evident in both zinc finger domains. The secondary chemical shifts in the linker between secondary structures are positive, but smaller in magnitude than for the zinc finger domains, suggesting that the helical conformation of the linker is not fully populated (and dynamically averaged). This observation is consistent with elevated values of *J*(0.87ω_H_) that were observed for the linker compared with the zinc finger domains (not shown). **(f, g) Secondary structural elements from backbone chemical shifts for MmGtsf1-115.** Helical (red) and strand (purple) backbone conformations were determined from assigned backbone chemical shifts using (f) TALOS and (g) CSI methods. Shaded boxes highlight the regions corresponding to the two zinc finger domains.

**Supplemental Figure 3.**
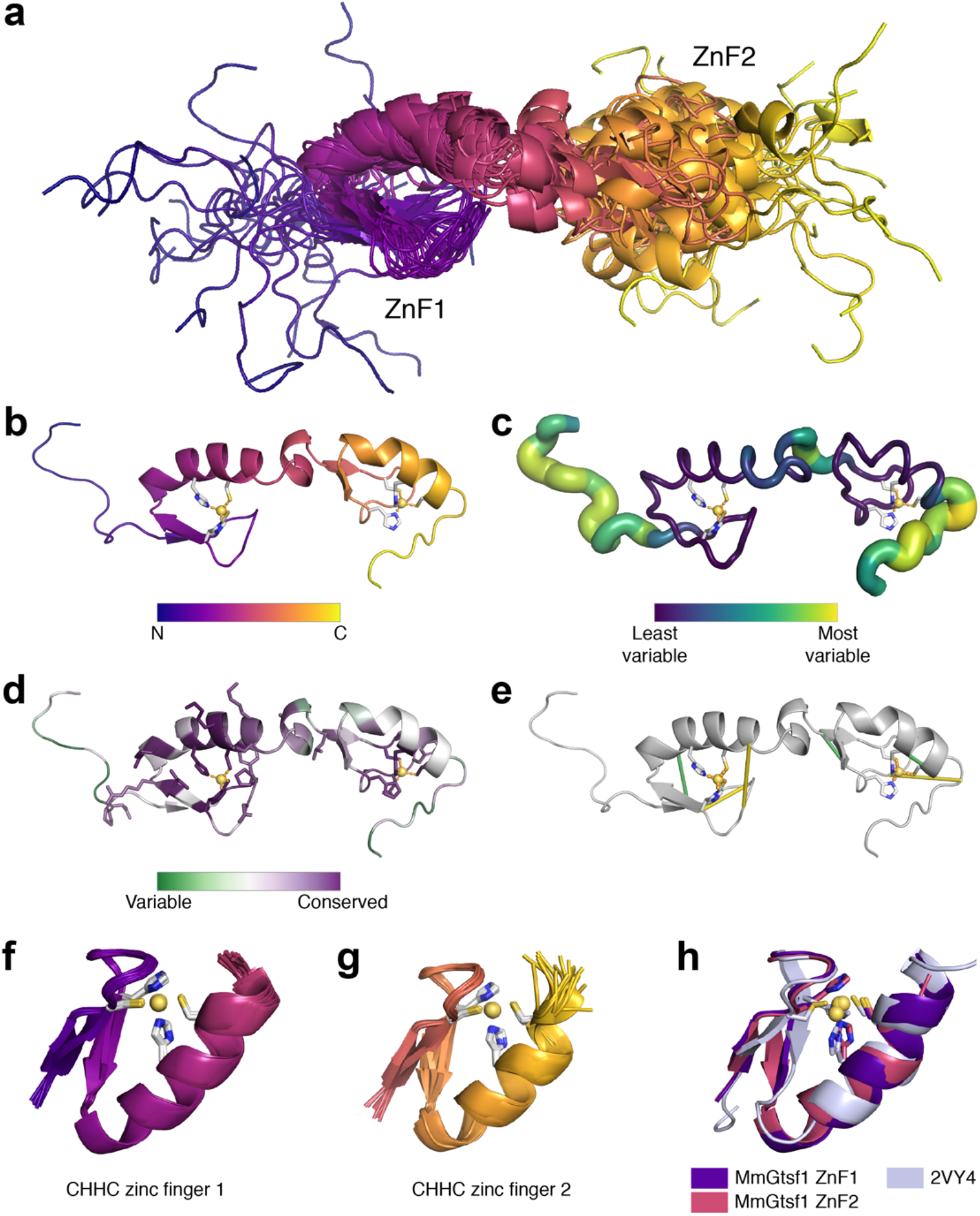
Structural analysis of MmGtsf1. **(a) NMR structural ensemble.** The 20 lowest energy structures of MmGtsf1 (residues 1-80) are shown as ribbon diagrams. Each display two tandem CHHC zinc fingers (ZnF1, ZnF2) with a flexible helix-containing linker. **(b) Single structure of MmGtsf1.** The lowest energy structure, presented as a ribbon, coloured as in (a). **(c) Local variability analysis.** Regions with greater local variability are indicated with a larger ribbon radius in addition to colour coding. The N-terminal, C-terminal, and linker regions showed the most variation. **(d) Conservation analysis.** In addition to the invariant, zinc-coordinating, CHHC residues, other highly conserved regions map to the positively-charged RNA-binding surface of ZnF1 (sticks). **(e) Co-evolution analysis.** Several intra-ZnF residues in special proximity to each other showed evidence of co-evolution (for example L15 and F30, with Van der Waals interactions between β-strand 1 and α-helix 1 of ZnF1). **(f, g) Superposition of individual zinc finger domains.** Structural alignments of the 20 lowest energy structures for (f) ZnF1 and (g) ZnF2 illustrate that the individual zinc fingers are consistently determined. **(h) Comparison of CHHC Zinc Finger Structures.** A total of three CHHC zinc finger structures have been determined to date (two from this work and another from the U11-48K protein of the minor spliceosome; PDB 2VY4). Each displays a strand-strand-helix architecture. For panels b-h, zinc-coordinating residues are shown as sticks. Zinc atoms are shown as yellow spheres.

**Supplemental Figure 4.**
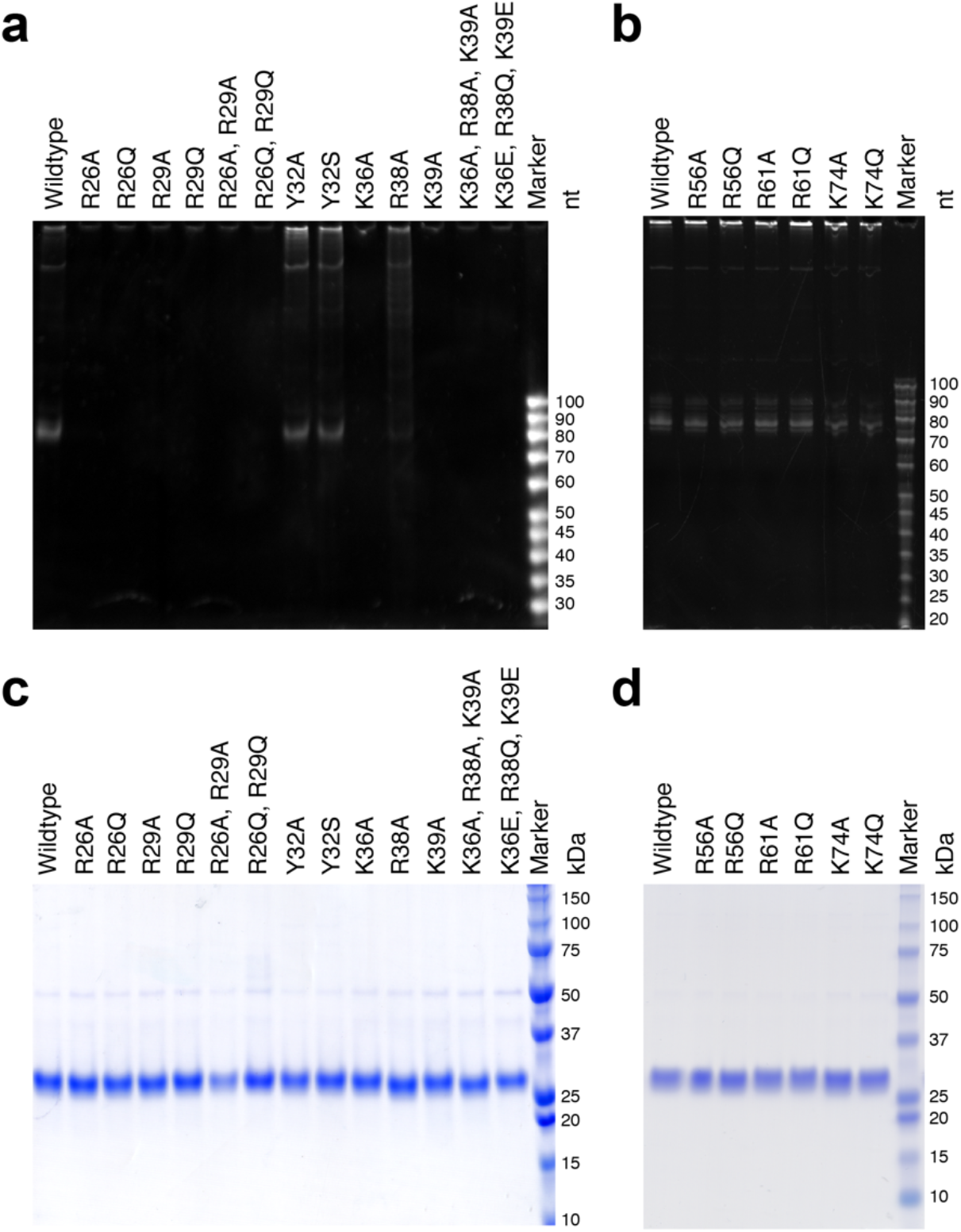
Point-mutagenesis analysis of MmGtsf1 on RNA-binding. **(a, b) Urea-PAGE assessment of the presence and size of RNAs co-purifying with MmGtsf1 constructs expressed in** *Sf9*. (a) Mutation of certain basic residues in ZnF1 abrogate RNA-binding activity including R26, A29, K36, and K39. The wildtype protein, which binds RNAs in the 70-90 nucleotide size range is shown for comparison in the first lane. (b) Basic residues in ZnF2 do not demonstrate significant effects on RNA pulldown. Each Urea-PAGE lane was loaded with a consistent amount of purified protein (1 μg). **(c, d) SDS-PAGE of affinity-purified MmGtsf1 constructs.** All of the tested (c) ZnF1 and (d) ZnF2 mutants could be expressed and purified at similar levels.

**Supplemental Figure 5.**
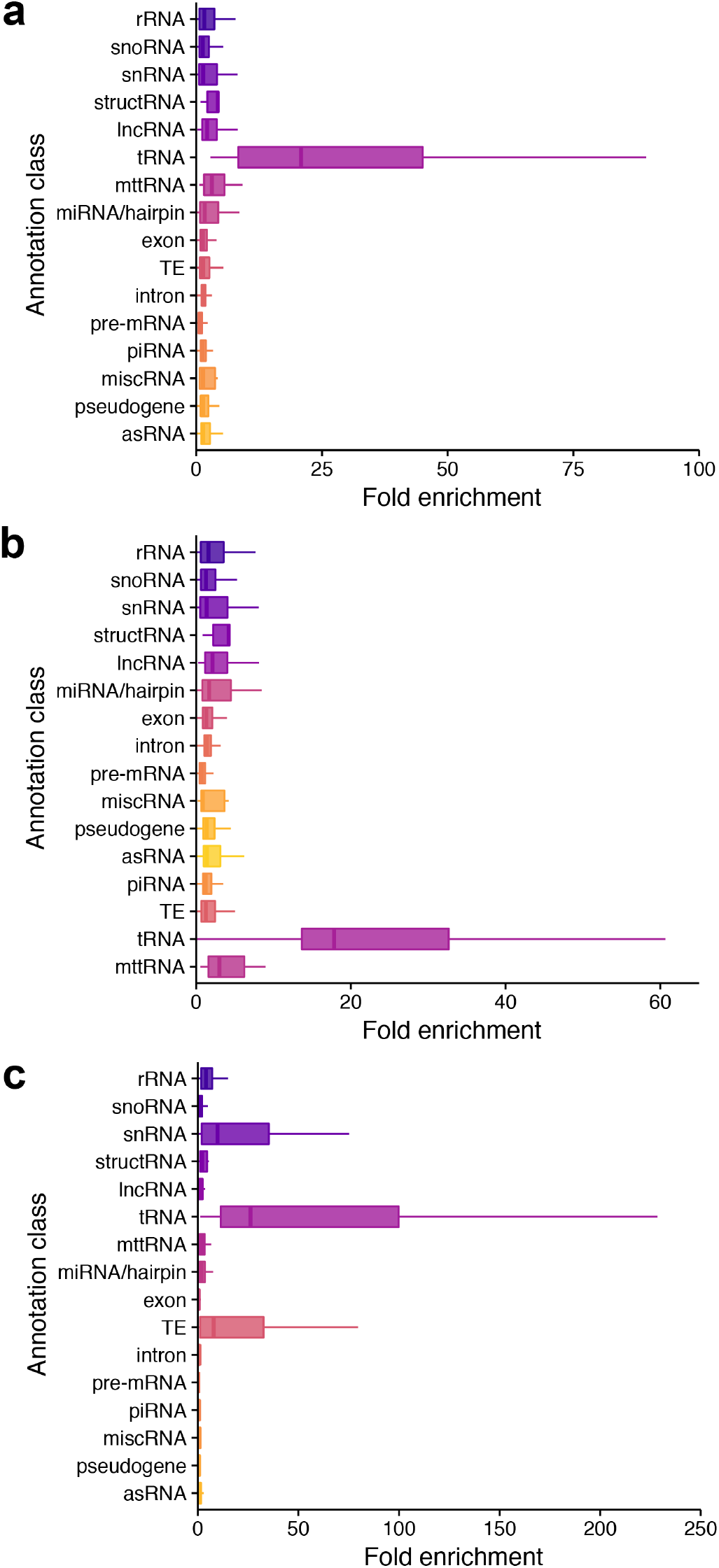
Comparison of Informatics Workflows – eCLIP for MmGtsf1-TEV-Strep in P19 cells. Each boxplot displays fold enrichment distributions for each gene annotation class. For convenience in comparison, panel (a) is duplicated from Fig. 1. Filtering order for each plot corresponds to the ordering of the annotation classes listed, from top to bottom. **(a) Analysis with the standard workflow based on abundance.** The standard workflow assigns read annotations with preference to annotation classes known to be highly abundant (e.g. rRNAs, snRNAs, and tRNA). **(b) Stringent analysis.** This workflow assigns read annotations such that the classes of interest in this study (piRNAs, transposable elements, and tRNAs) are assigned last. This results in fewer reads being assigned in these classes, but with those reads being less ambiguous in their source. **(c) Analysis using an alternative gene model.** Abundance-based filtering was performed as in (a), but employing an alternative gene model to track multi-mapping reads. This analysis allowed for standardized gene expression analysis tools to be used, in this case, DESeq2. In each panel, the filtering order is indicated by the order of annotations on the y-axis, with the upper-most annotations getting the highest priority. Colour-coding is consistent among panes for each annotation class. In each case, tRNAs exhibited a distribution with the greatest mean fold enrichment.

**Supplemental Figure 6.**
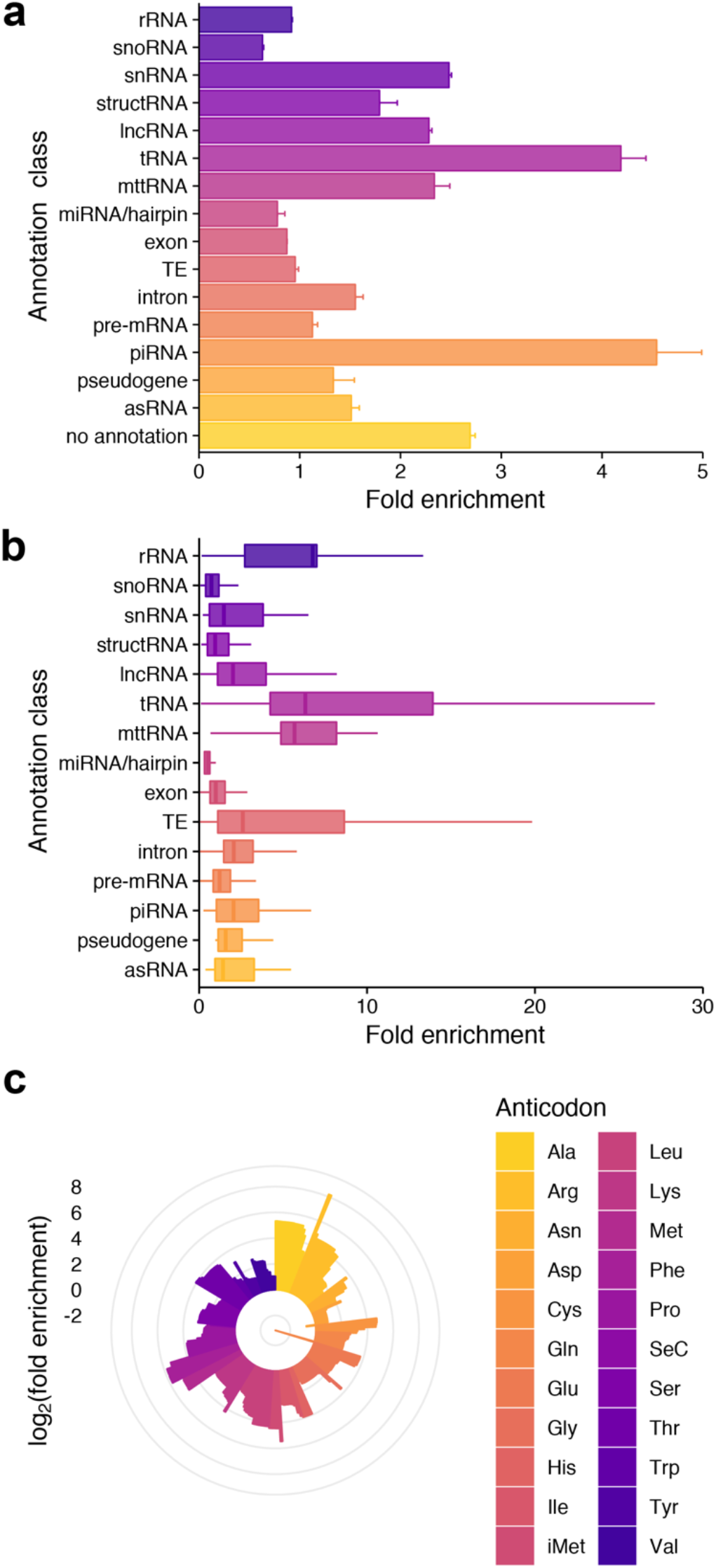
Informatics summary – eCLIP for FLAG-tagged Asterix in OSS cells (endogenous locus; tagged with CRISPR). **(a) Fold enrichment of reads mapping to each annotation class in the eCLIP pull-downs.** Error bars represent the standard error. **(b) Box-plot of fold enrichments for each gene within an annotation class.** Outliers are not shown. The colour coding for each class is consistent with the *Drosophila* data presented in Figure 2. The order of annotation classes in each panel (from top to bottom) indicates the order in which reads were filtered before being ascribed to subsequent classes. All eCLIP experiments were performed in duplicate. (c) **Fold enrichment calculated for each tRNA.** Colour coding corresponds to a given anti-codon. For panels a and b, filtering order corresponds to the ordering of the annotation classes listed, from top to bottom.

**Supplemental Figure 7.**
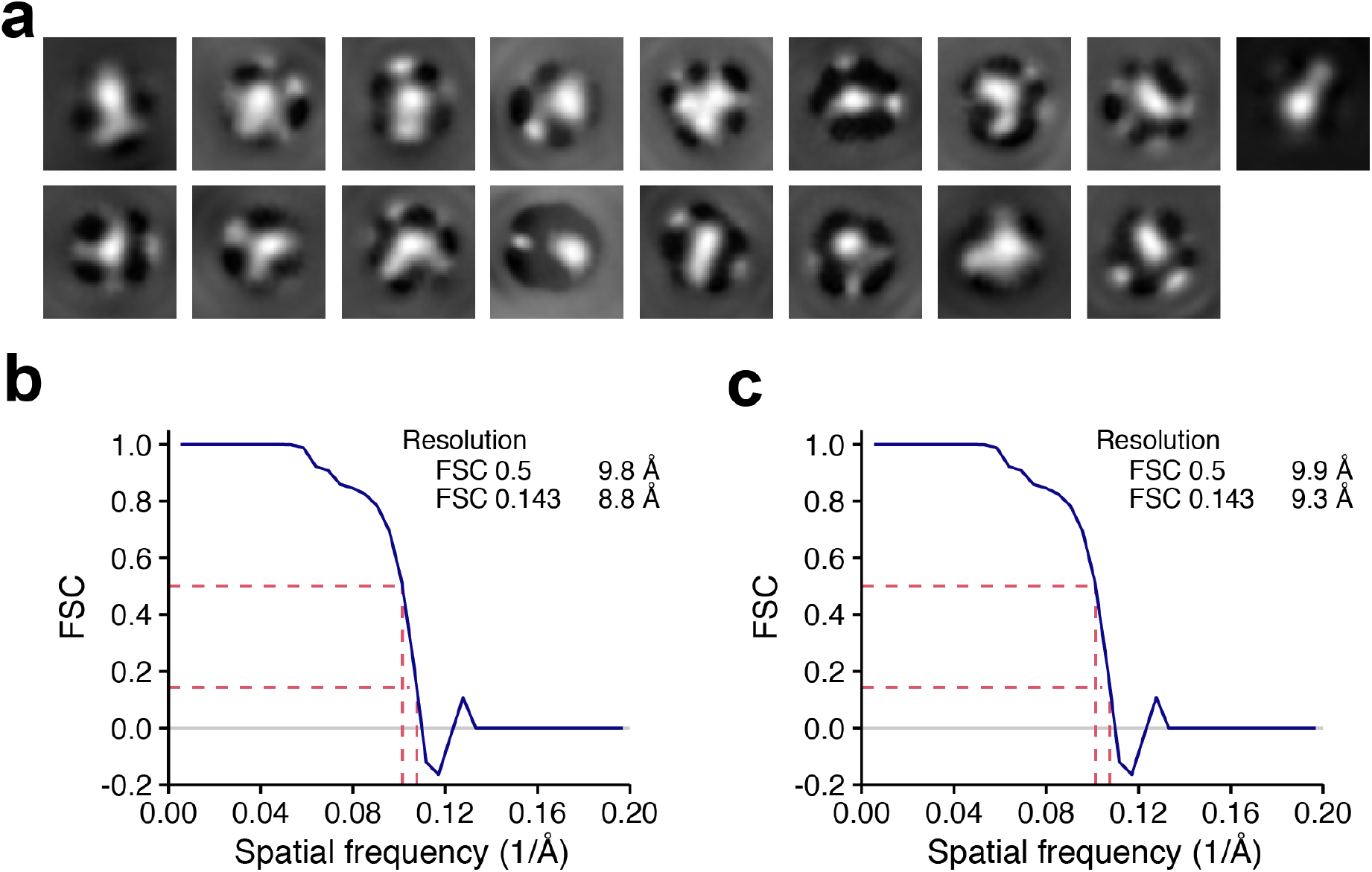
Cryo-electron microscopy structure determination of MmGtsf1 bound to co-purifying tRNA. (a) 2D class averages. Classification was performed with cisTEM and resulted in a number of averages where tRNA-like forms were readily observed. The class averages shown are for the full-length construct. **(b,c) FSC curves.** FSC curves corresponding to the reconstructions of (b) the full-length MmGtsf1 construct and (c) the MmGtsf1 truncation construct (only the first zinc finger; residues 1-45) in the presence of co-purifying RNA. Based on gold standard FSC, the structures have an estimated resolution of approximately 9 Å. However, despite the calculated values, visual inspection of the reconstructions suggests resolutions of 15-20 Å.

**Supplemental Figure 8.**
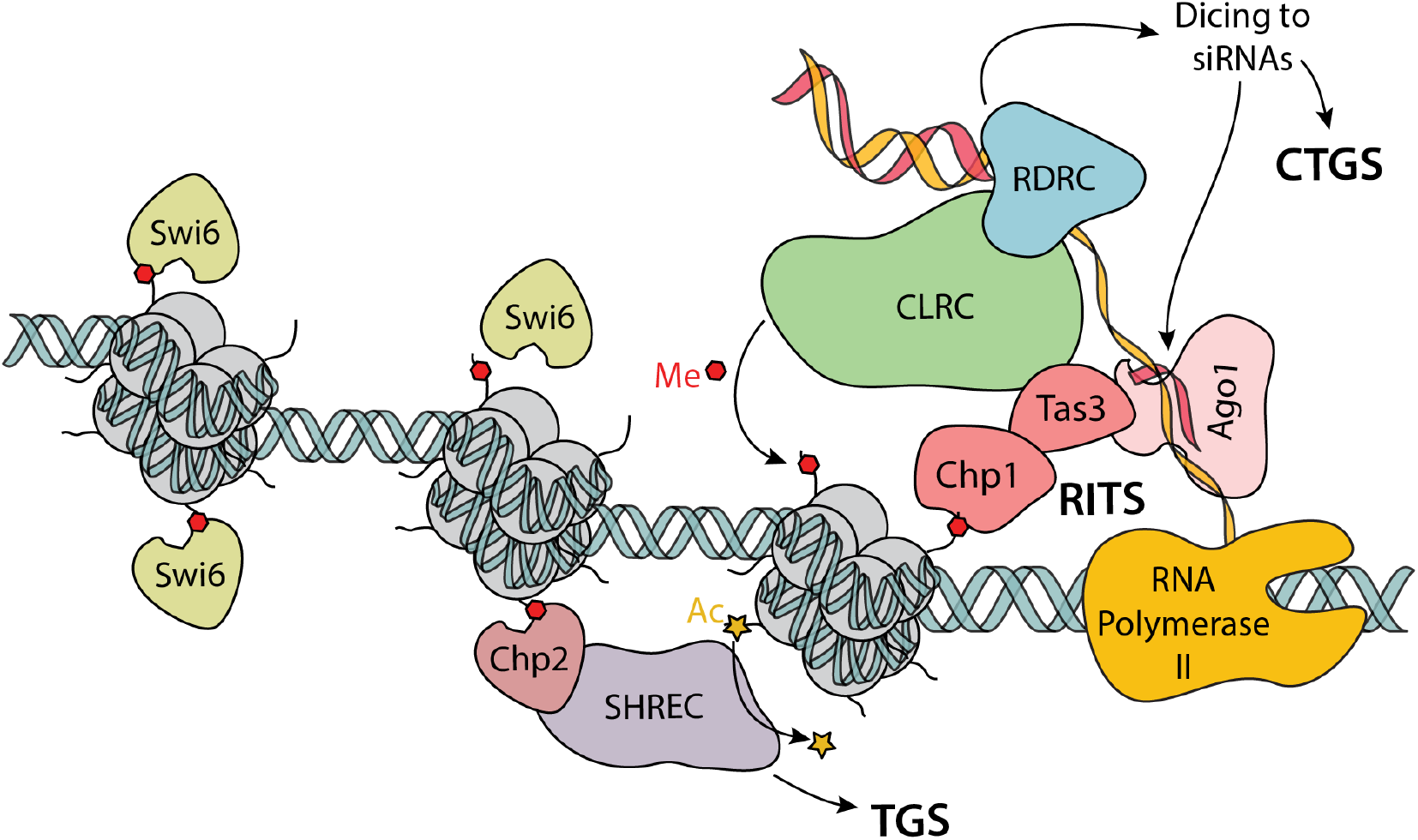
Diagram of similar silencing mechanisms in *S. pombe*. In yeast, multivalent interactions enforce silencing. Nascent transcripts (yellow) are recognized by guide-loaded Ago1 (pink) as well as RNA-directed RNA Polymerase Complex (RDRC; blue). In turn, the RITS complex, consisting of Chp1 and Tas3 (red), and CLRC (green) are able to methylate target loci. Additional silencing machinery, including Swi6 (beige) induce chromatin formation and lead to silencing. (CTGS = co-transcriptional gene silencing; RITS = RNAi-induced transcriptional silencing complex; TGS = transcriptional gene silencing; SHREC = Snf2/HDAC-containing Repressor Complex)

**Supplemental Table 1.** NMR and Refinement Statistics

| MmGtsf1 |  |  |
| --- | --- | --- |
| NMR distance and dihedral constraints |  |  |
| Distance constraints |  |  |
| Total NOE | 1022 |  |
| Intra-residue | 552 |  |
| Inter-residue | 470 |  |
| Sequential ($\vert i-j\vert = 1$) | 255 | |
| Medium-range ($\vert i-j\vert < 4$) | 99 | |
| Long-range ($\vert i-j\vert > 5$) | 116 | |
| Hydrogen bonds | 21 |  |
| Total dihedral angle restraints | 126 |  |
| $\phi$ | 46 | |
| $\psi$ | 48 | |
| Structure statistics |  |  |
| Number of models | 20 |  |
| Violations (mean and s.d.) |  |  |
| Distance constraints ($>0.1 \text{ \AA}$) | $1.71 \times 10^{-2} \pm 3.50 \times 10^{-3}$ | |
| Dihedral angle constraints ($^{\circ}$) | $0.3686 \pm 0.05$ | |
| Max. dihedral angle violation ($^{\circ}$) | $1.0 \pm 0.19$ | |
| Max. distance constraint violation ($\text{\AA}$) | $0.28 \pm 0.07$ | |
| Deviations from idealized geometry |  |  |
| Bond lengths ($\text{\AA}$) | $2.97 \times 10^{-3} \pm 7.27 \times 10^{-5}$ | |
| Bond angles ($^{\circ}$) | $0.44 \pm 9.84 \times 10^{-3}$ | |
| Impropers ($^{\circ}$) | $1.12 \pm 8.27 \times 10^{-2}$ | |
| Average pairwise r.m.s. deviation ($\text{\AA}$) | CHHC Zinc Finger 1<br>Residues (14-38) | CHHC Zinc Finger 2<br>Residues (48-68) |
| Heavy | $0.94 \pm 0.12$ | $0.91 \pm 0.10$ |
| Backbone | $0.22 \pm 0.04$ | $0.22 \pm 0.04$ |
| Validation | Full Structure<br>(Residues 1-80) | Core<br>(Residues 13-73) |
| Ramachandran favored | $71.1 \pm 0.5$<br>($91.15\% \pm 0.64\%$) | $56.75 \pm 0.24$<br>($96.19\% \pm 0.41\%$) |
| Ramachandran outliers | $0.75 \pm 0.18$<br>($0.96\% \pm 0.23\%$) | $0.05 \pm 0.05$<br>($0.08\% \pm 0.08\%$) |
| Favored rotamers | $69.25 \pm 0.42$<br>($92.33\% \pm 0.55\%$) | $51.55 \pm 0.37$<br>($92.05\% \pm 0.65\%$) |
| Poor rotamers | $0.4 \pm 0.17$<br>($0.53\% \pm 0.22\%$) | $0.25 \pm 0.14$<br>($0.45\% \pm 0.25\%$) |
| Molprobrity Score | $2.28 \pm 0.04$ | $1.95 \pm 0.05$ |
| C$_{\beta}$ deviations | $0.0 \pm 0.0$ | $0.0 \pm 0.0$ |
| Bad bonds | $0.0 \pm 0.0$ | $0.0 \pm 0.0$ |
| Bad angles | $0.0 \pm 0.0$ | $0.0 \pm 0.0$ |
| Cis prolines | $0.0 \pm 0.0$ | $0.0 \pm 0.0$ |
| C$_{\alpha}$ geometry outliers | $0.1 \pm 0.07$ | $0.0 \pm 0.0$ |

**Supplemental Table 2.**
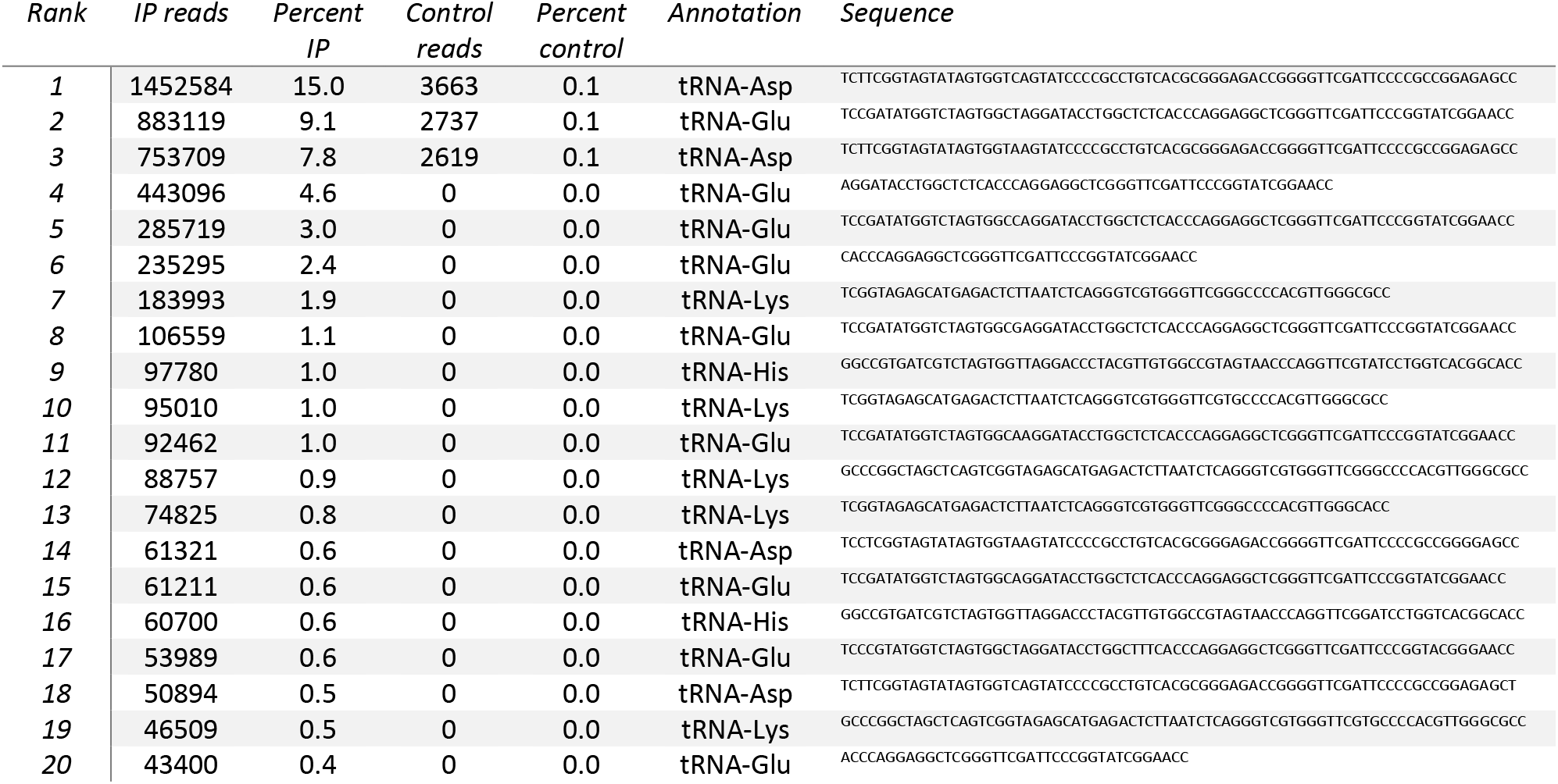
Top 20 most abundant reads determined by RNA sequencing from MmGtsf1-TEV-Strep recombinant over-expression in *Sf9*, followed by pull-down. tRNAs made up a very substantial part of the library, with approximately 50% of all reads corresponding to these top 20 sequences. Similar results were found for replicate library preparations.

